# Dissecting the regulatory logic of specification and differentiation during vertebrate embryogenesis

**DOI:** 10.1101/2024.08.27.609971

**Authors:** Jialin Liu, Sebastian M. Castillo-Hair, Lucia Y. Du, Yiqun Wang, Adam N. Carte, Mariona Colomer-Rosell, Christopher Yin, Georg Seelig, Alexander F. Schier

**Author notes:** Equal contribution: these authors contributed equally and are listed alphabetically (Sebastian M. Castillo-Hair, Lucia Y. Du).

## Abstract

The interplay between transcription factors and chromatin accessibility regulates cell type diversification during vertebrate embryogenesis. To systematically decipher the gene regulatory logic guiding this process, we generated a single-cell multi-omics atlas of RNA expression and chromatin accessibility during early zebrafish embryogenesis. We developed a deep learning model to predict chromatin accessibility based on DNA sequence and found that a small number of transcription factors underlie cell-type-specific chromatin landscapes. While Nanog is well-established in promoting pluripotency, we discovered a new function in priming the enhancer accessibility of mesendodermal genes. In addition to the classical stepwise mode of differentiation, we describe instant differentiation, where pluripotent cells skip intermediate fate transitions and terminally differentiate. Reconstruction of gene regulatory interactions reveals that this process is driven by a shallow network in which maternally deposited regulators activate a small set of transcription factors that co-regulate hundreds of differentiation genes. Notably, misexpression of these transcription factors in pluripotent cells is sufficient to ectopically activate their targets. This study provides a rich resource for analyzing embryonic gene regulation and reveals the regulatory logic of instant differentiation.

## Main

Cell differentiation can generally be conceptualized as a stepwise process. It starts with specification to a developmental fate and progresses through sequential changes in gene expression and morphology to ultimately result in the acquisition of specialized structures and functions (Liberali & Schier, 2024). For example, insulin-producing beta cells derive from a stepwise specialization through pluripotent, endodermal, pancreatic and endocrine states (Murtaugh, 2007). However, certain cell types rapidly differentiate during early embryogenesis without undergoing multi-step transitions These include the trophectoderm in mammals (Lim et al., 2020), and the liver-like yolk syncytial layer (YSL) and the skin-like enveloping layer (EVL) in fish (Kimmel et al., 1990, 1995). This phenomenon of ‘instant differentiation’ is characterized by its speed and simplicity compared to stepwise differentiation. For example, at the onset of gastrulation, the EVL has already differentiated, displaying distinct morphology (Kimmel et al., 1990), cytoskeleton and adhesion structures (Zalik et al., 1999), with hundreds of genes differentially expressed compared to embryonic cells (Satija et al., 2015). Although this phenomenon has long been described, the gene regulatory logic of instant differentiation has remained elusive.

Recent advances to systematically study gene regulatory networks (GRNs) provide the opportunity to address this question (Badia-i-Mompel et al., 2023; Fleck et al., 2023; Janssens et al., 2022; Kamimoto et al., 2023; Saunders et al., 2023). For example, single-cell RNA sequencing (scRNA-seq) has led to the systematic reconstruction of the gene expression trajectories during cell type specification and differentiation (Briggs et al., 2018; Farrell et al., 2018; Qiu et al., 2022; Wagner et al., 2018). More recently, these efforts have been complemented with single-cell measurements of chromatin accessibility (Argelaguet et al., 2019; Cao et al., 2018; Ma et al., 2020). Deep learning models trained on these data can predict cell type accessibility from sequence and can be interrogated to reveal *cis*-regulatory elements (Ameen et al., 2022; Eraslan et al., 2019; Janssens et al., 2022; Zhou & Troyanskaya, 2015). Furthermore, the capacity to correlate changes in the epigenome with alterations in the transcriptome facilitates the construction GRNs (Badia-i-Mompel et al., 2023; Bravo González-Blas et al., 2023; Fleck et al., 2023). In this study, we created a single-cell multi-omics (scMultiome) atlas that jointly tracks RNA expression and chromatin accessibility throughout early zebrafish embryogenesis. We utilized this resource to develop a deep learning model and construct GRNs to dissect the gene regulatory logic of instant differentiation.

### Single-cell multi-omics identifies gene expression and chromatin accessibility dynamics during embryogenesis

Single-cell gene expression (Farrell et al., 2018; Wagner et al., 2018) or chromatin accessibility (Sun et al., 2024) have been profiled separately during zebrafish embryogenesis. However, we lack one-to-one cell correspondence and even stage correspondence between the two modalities, which would provide a more detailed and integrated understanding of cellular states and regulatory mechanisms. To systematically dissect the gene regulatory logic that governs embryogenesis, we simultaneously measured gene expression and chromatin accessibility within individual nuclei during early zebrafish embryogenesis (Figure 1 A, 10x Genomics). Our comprehensive dataset spans nine developmental stages, from pluripotency to early organogenesis (high stage at 3.3 hours post-fertilization (hpf) to 6-somite stage at 12 hpf). In total, 40,992 high-quality single nuclei were captured, with a median of 2,082 expressed genes and 16,925 ATAC fragments detected per nucleus (Table S1), an improvement to previous standalone single-cell RNA sequencing (scRNA-seq) (Farrell et al., 2018) and single-nucleus ATAC sequencing (snATAC-seq) (Sun et al., 2024) datasets. For quality control we generated standalone snATAC-seq data using the same 10x Genomics technology for 21,050 nuclei at the onset of gastrulation (50% epiboly) and 6-somite stage. The chromatin accessibility between the scMultiome and standalone snATAC-seq was highly concordant (Figure S1). Using the snRNA-seq modality of the scMultiome, we identified 95 cell states based on previously described cell state markers (Farrell et al., 2018; Qiu et al., 2022; Wagner et al., 2018). These cell states were also independently found in the snATAC-seq modality-based UMAP space, revealing that both modalities can capture cellular diversity (Figure 1A and Figure S2). These results indicate that the single-cell multi-omics data is of high quality.

**Figure 1:**
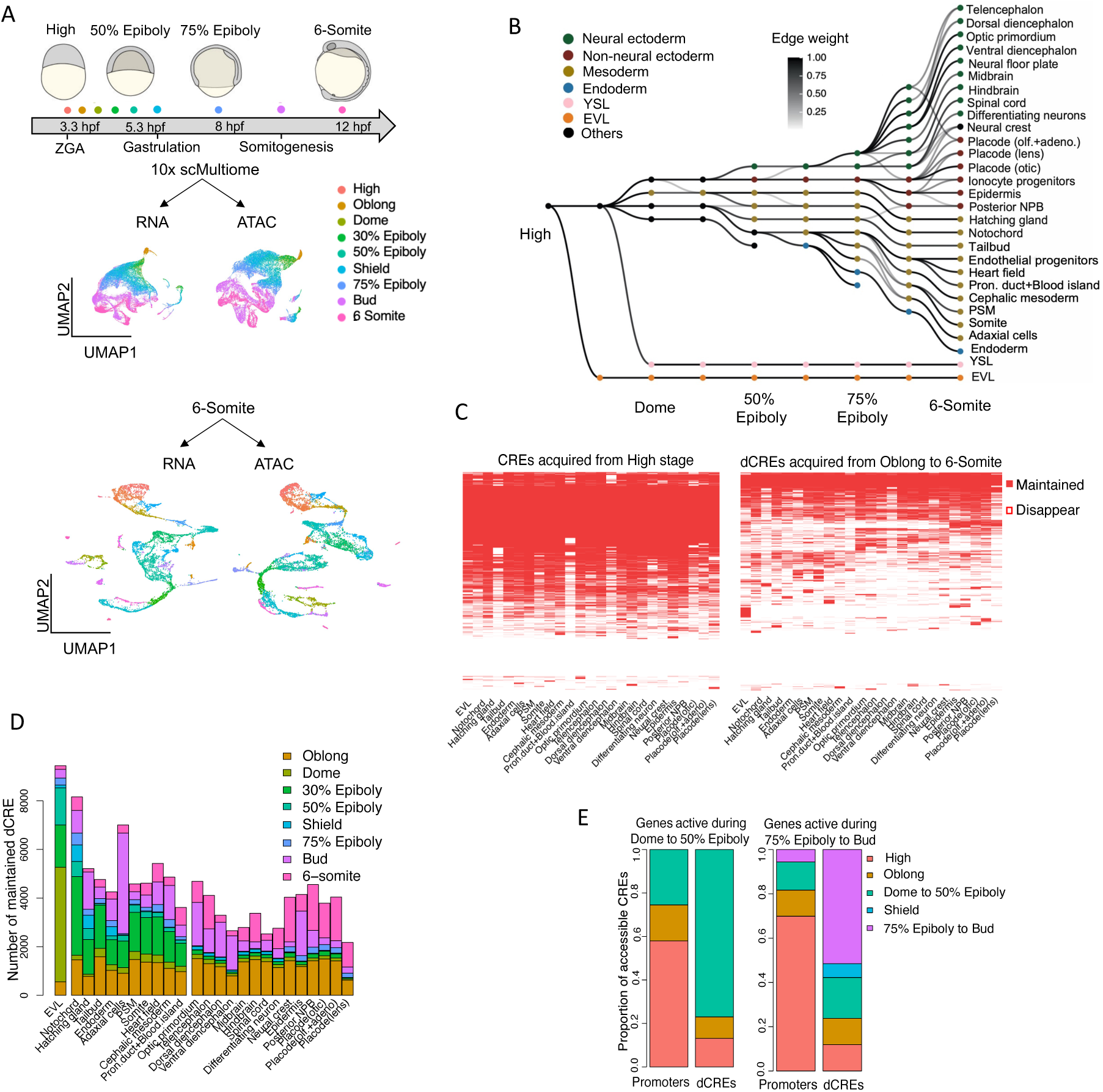
Zebrafish single-cell multi-omics atlas reveals gene expression and chromatin accessibility dynamics. A. Upper panel: single-cell multi-omics data were collected from zebrafish embryos at nine developmental stages (represented by colored dots) using 10x Genomics technology. Middle panel: Uniform Manifold Approximation and Projection (UMAP) visualization of single cells, with each cell colored according to its developmental stage. The UMAP coordinates are based on either RNA or ATAC-seq data. Lower panel: UMAP visualization of single cells at 6-somite stage, colored by clusters. The clusters are defined by gene expression, and the UMAP coordinates are derived from either RNA or ATAC-seq data. B. Inferred relationships between cell states during early zebrafish embryogenesis. Each row corresponds to a specific cell state, while each column represents a developmental stage. The nodes are color-coded according to different germ layers. All edges with weights above 0.2 are displayed. olf.+adeno., olfactory + adenohypophyseal; NPB, neural plate border; Pron., pronephros; PSM, presomitic mesoderm; EVL, enveloping layer; YSL, yolk syncytial layer. C. Analysis of distal cis-regulatory elements (dCREs) acquired at high stage (left panel) or any stage from oblong to 6-somite (right panel), examining which elements are maintained at or disappear by the 6-somite stage. Note that most dCREs at high stage are maintained in most cell types throughout early embryogenesis, whereas dCREs acquired later disappear or become cell-type-specific. D. The number of acquired dCREs at each stage that are maintained by the 6-somite stage. The figure is divided into three groups: EVL, mesendoderm, and ectoderm, shown from left to right. Note that the timing of CRE appearance generally coincides with the timing of cell type specification or differentiation. E. The proportion of promoters or dCREs that are accessible at each stage. Note that promoters are often accessible before the expression of the corresponding gene.

To characterize the chromatin accessibility profiles of each cell state, we used a cluster-specific and replicate-aware peak calling approach (Granja et al., 2021). We identified 444,530 peaks, representing putative *cis*-regulatory elements (CREs). Distal CREs (dCREs), including putative enhancers, were defined as peaks >500 bp from an annotated transcription start site (TSS). Peaks within <500 bp of a TSS were annotated as promoters. To link cell states and types across different stages into trajectories, we used gene expression similarity between adjacent stages(Briggs et al., 2018; Qiu et al., 2022) (Figure 1B, Table S2). Briefly, we connected each cell in a given stage to its most likely ancestors in the preceding stage based on Euclidean distances in a low-dimensional space (see methods). The reconstructed trajectories recapitulated the developmental paths described in previous studies (Farrell et al., 2018; Qiu et al., 2022; Wagner et al., 2018). The earliest branching events include the segregation between embryonic and extra-embryonic cell types; i.e. the enveloping layer (EVL) and the yolk syncytial layer (YSL). Early differentiation of EVL and YSL provides epidermal and lipid-metabolizing functions, respectively, which are essential for embryo development. From 50% epiboly (onset of gastrulation), we observed the expansion of mesendodermal cell types, while ectodermal cell types expand from the bud stage (end of gastrulation), culminating in the presence of two dozen cell types at 6-somite stage, including heart, adaxial muscle, forebrain, epidermis, hatching gland, and notochord. In summary, the scMultiome dataset provides an extensive resource to explore chromatin accessibility and gene expression during early zebrafish embryogenesis. In the following sections we highlight how the scMultiome dataset can provide new biological insights.

### Chromatin accessibility dynamics reveal cell-type- and stage-specific cis-regulatory element usage

To analyze the temporal dynamics of putative distal cis-regulatory element (dCREs) usage, we asked how many dCREs acquired at a given stage were still present at 6-somite stage (Figure 1C and 1D). We found that more than 50% of the dCREs acquired at the high stage were maintained at the 6-somite stage. These dCREs were ubiquitously accessible in all cell types (Figure 1C) and associated with housekeeping genes (e.g. genes involved in metabolic processes and RNA biosynthetic processes) (Table S3). In contrast to these ubiquitous dCREs, dCREs acquired and maintained after the high stage were mostly cell-type-specific (Figure 1C). The timing at which these cell-type-specific dCREs are acquired coincided with the specification and differentiation of the associated cell types (Figure 1D and Figure S3). Three waves were apparent: EVL and YSL (Figure S3) dCREs at the 6-somite originate from the oblong to the 50% epiboly stages (from blastula to the onset of gastrulation); mesoderm-associated dCREs appear at 30% epiboly (before the onset gastrulation); dCREs in ectoderm-derived cell types emerge from the bud (end of gastrulation) to the 6-somite stage. These results reveal a hierarchical appearance of dCREs, in which ubiquitous dCREs are acquired early, whereas cell-type-specific dCREs appear during cellular specification and differentiation.

In addition to studying the relationship of dCRE usage with cell type diversification, the simultaneously measured two modalities allow us to compare the temporal emergence of CREs with the timing of gene activation. We identified cell-type-specific genes that become active from dome to 50% epiboly (onset of gastrulation). We inferred their associated dCREs (association score>0.1 and p-value<0.05) based on the correlation analysis between chromatin accessibility and gene expression (see methods). We observed that the majority of dCREs become accessible at the time when gene expression starts (Figure 1E). In contrast, promoter peaks generally preceded gene expression. For example, for genes active during 75% epiboly to bud stage (mid- to end of gastrulation), the majority of promoters are already accessible at the high stage (Figure 1E). These results suggest that promoters are generally primed for future gene expression (Pálfy et al., 2020; Reddington et al., 2020), whereas the accessibility of most dCREs coincides with gene activation.

### Nanog primes chromatin accessibility of putative mesendodermal distal *cis*-regulatory elements

While most putative dCREs become accessible when gene expression starts, some dCREs are accessible beforehand (Figure 1E), reminiscent of the concept of enhancer priming (Spitz & Furlong, 2012). In this process, transcription factors (TFs) bind to an enhancer earlier in embryogenesis, rendering it accessible for binding by subsequent TFs that activate gene expression during later stages. Enhancer priming has been suggested to enable rapid and sustained transcriptional responses (Falo-Sanjuan et al., 2019). Multi-omics profiling during mouse embryogenesis indicated that neuroectodermal but not mesendodermal enhancers are primed by chromatin accessibility in the epiblast (Argelaguet et al., 2019). To investigate if this property is conserved in zebrafish, we analyzed if and when dCREs associated with mesendodermal and neuroectodermal genes are primed. In contrast to mouse embryogenesis, we found that 13% of dCREs associated with mesendodermal genes active between dome and 50% epiboly are already accessible at the high stage (Figure 2A and 2B) when most cells are pluripotent. This percentage is significantly higher than that of non-associated dCREs (6%). In contrast to the mesendoderm, we did not find a higher proportion of high-stage accessible dCREs associated with neuroectodermal genes (active from 75% epiboly to the bud stage, the end of gastrulation) (Figure 2C). Instead, at 50% epiboly (onset of gastrulation), the proportion of accessible neuroectodermal associated dCREs is significantly higher than that of non-associated dCREs (Figure 2C and D). These results suggest that dCRE priming occurs in both mesendoderm and neuroectoderm and that the timing of priming correlates with the temporal specification of these tissue types.

**Figure 2:**
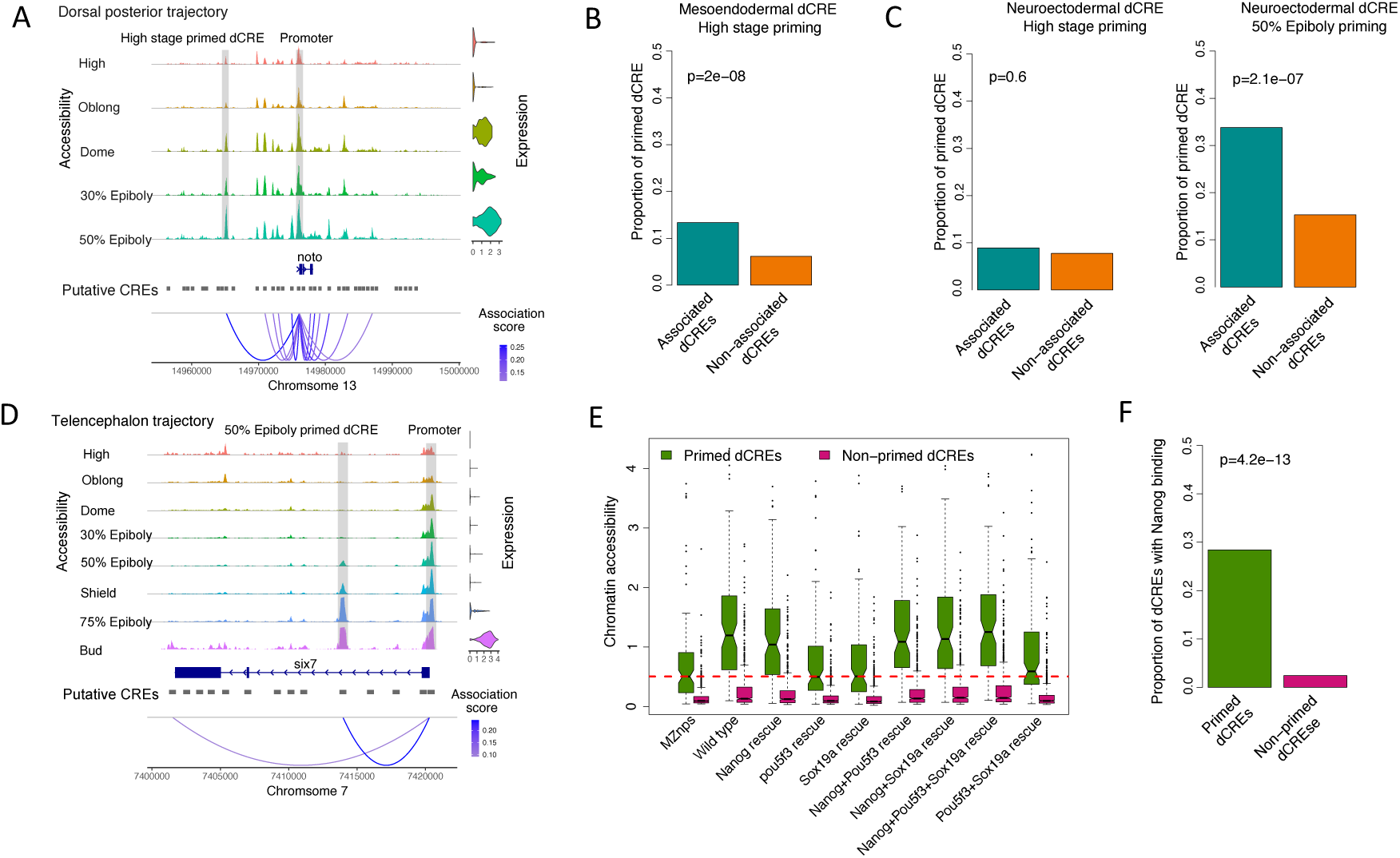
Nanog primes putative mesendodermal distal *cis*-regulatory elements. A. Left panel: representative chromatin accessibility tracks for all CREs near TSS of the *noto* gene across the dorsal mesoderm trajectory. The aggregated pseudo-bulk data for chromatin accessibility in each cell state is presented. Right panel: violin plot shows the expression of *noto* across the dorsal mesoderm trajectory. Bottom panel: the numbers above chromosome 13 indicates the genomic coordinates. B. Proportion of mesendodermal putative distal *cis*-regulatory elements (dCREs) primed at the high stage. For each mesendodermal gene, we inferred its associated dCREs from all dCREs located within 50 kb upstream or downstream of the transcription start site (TSS). dCREs were considered associated if they had an association score >0.1 and a p-value <0.05. “High stage priming” indicates that these dCREs were already accessible at the high stage. C. The left bar graph shows the proportion of neuroectodermal dCREs that were primed at the high stage. The right bar graph illustrates the proportion of neuroectodermal dCREs that were primed at the 50% epiboly stage. Similar to mesendodermal genes, for each neuroectodermal gene we inferred its associated dCREs. “High stage priming” means these dCREs were accessible at the high stage, while “50% Epiboly priming” means they were accessible at the 50% epiboly stage. D. Similar to A, but for telencephalon trajectory. E. Chromatin accessibility of primed versus non-primed dCREs across various rescue conditions. Here, the dCREs only refer to the associated ones. The lower and upper intervals indicated by the dashed lines (“whiskers”) represent 1.5 times the interquartile range (IQR), and the box shows the lower and upper intervals of IQR together with the median. F. The proportion of primed versus non-primed dCREs that overlap with Nanog peaks at the high stage. Here, the dCRES only refer to the associated ones. The p-values for B, C and F were determined using pairwise Fisher’s exact tests.

To investigate which TFs prime mesendodermal dCREs at high stage, we focused on three maternally deposited pioneer TFs, Nanog, Pou5f3 and Sox19b. We compared the chromatin accessibility of primed mesendodermal dCREs between triple mutant and various rescue conditions, using previously published data (Miao et al., 2022). First, non-primed dCREs showed much lower accessibility than primed dCRE (Figure 2E). Second, we discovered that chromatin accessibility increases whenever the rescue includes Nanog, with the greatest increase observed in the triple rescue. In contrast, the rescue conditions with just Pou5f3 and Sox19b contributed little to chromatin accessibility. These observations suggest that Nanog is the primary factor that primes mesendodermal dCREs. To further test this idea, we examined the overlap between primed mesendodermal dCREs and Nanogbinding sites using CUT&RUN data from the high stage (X. Wang et al., 2022). We found that approximately 30% of primed mesendodermal dCREs overlap with NanogCUT&RUN peaks, a proportion much higher than the overlap observed with non-primed mesendodermal dCREs (Figure 2F). Overall, these results indicate that the dCREs of numerous mesendodermal genes are primed primarily by the pioneer TF Nanog.

### Deep learning identifies a small number of transcription factors that predict cell type-specific chromatin landscapes

Deep-learning models, such as convolutional neural networks (CNNs), have emerged as powerful tools for predicting genomic “activity profiles” based on DNA sequences (Eraslan et al., 2019). Interpretative methods (Novakovsky et al., 2022), can extract sequence motifs responsible for the predicted “activity profiles” and help decode the *cis*-regulatory logic of gene expression. To systematically investigate the sequence motifs of chromatin accessibility, we developed DeepDanio, a deep learning model (Figure 3A, see methods). DeepDanio was trained to predict chromatin accessibility in all 95 cell states across all developmental stages given a 500bp DNA sequence. To improve accuracy and generalization, we designed DeepDanio as an ensemble of three deep CNNs with residual connections (Figure 3A and Table S7), where each CNN was trained on CREs from a different subset of chromosomes (∼80% of all CREs), with the remaining CREs used for early stopping to prevent overfitting (2 chromosomes, 10% of all CREs) and for performance evaluation (2 chromosomes, 10% CREs). On sequences held out from training, DeepDanio showed good performance in predicting accessibility (Figure 3B). Overall, 92.4% of test CREs were predicted with a statistically significant correlation coefficient (Figure 3C, permutation test, p-value <0.01). Additionally, across all CREs within each specific cell state, DeepDanio attained a high correlation between observed and predicted chromatin accessibility (Figure S4A).

**Figure 3:**
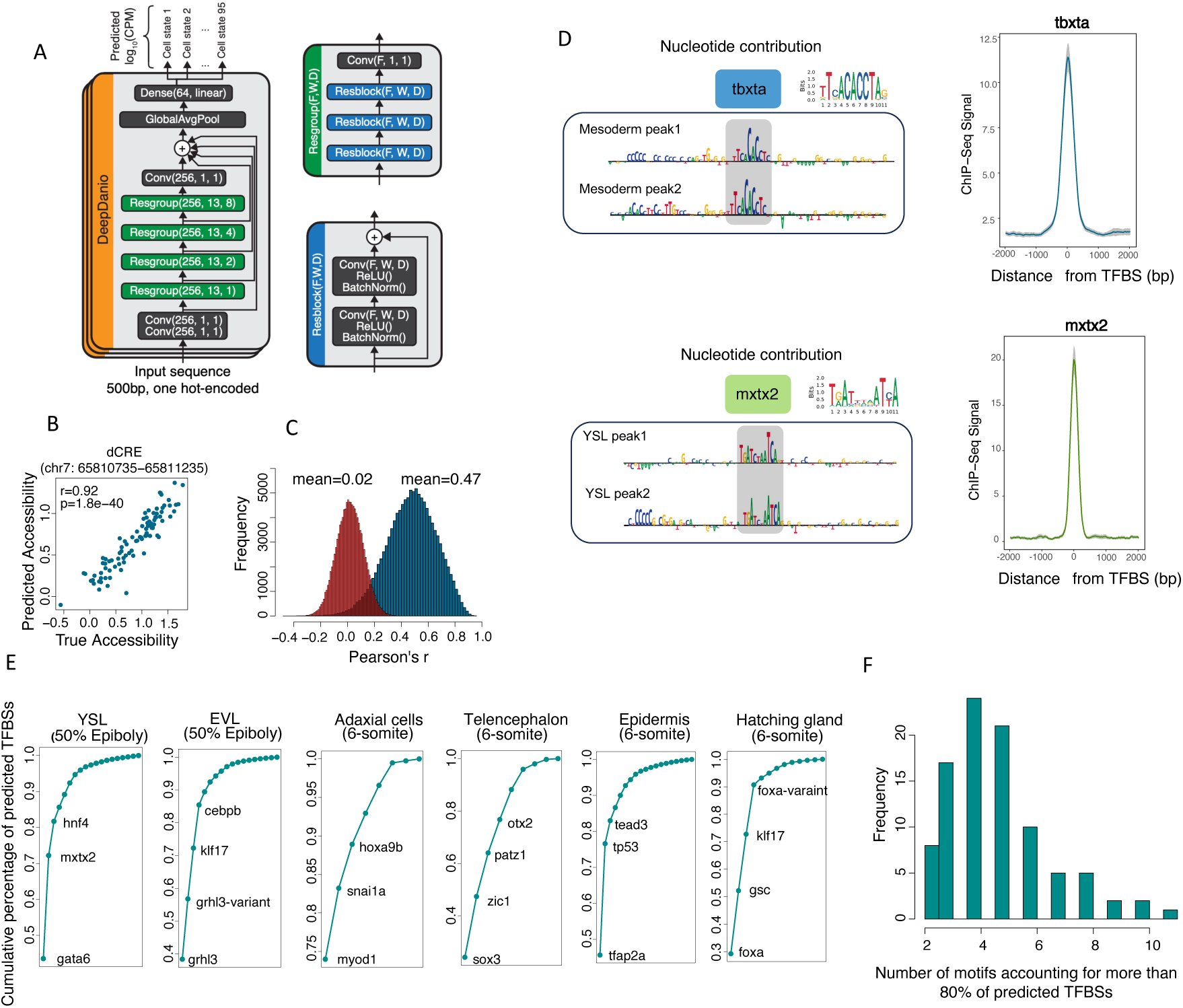
DeepDanio predicts chromatin accessibility based on DNA sequence and identifies sequence motifs of chromatin accessibility. A. Architecture of DeepDanio. In each model, a 500bp sequence is processed via a series of residual blocks containing convolutional layers and skip connections. The model output is the predicted log10(CPM) (accessibility) of each of the 95 cell states, obtained as the average of three independently trained models. B. Measured vs. predicted accessibility for all cell states on a selected CRE held out from training, along with their Pearson correlation coefficient (r) and p-value. C. Distribution of Pearson correlation values across all CREs held out from training (blue). CREs in the evaluation set of all three individual models and their predictions were pooled together. To assess the statistical significance of the observed correlation, a null distribution was generated by randomly shuffling the cell type labels of the predictions (red, permutation test). The p-value was inferred by comparing the observed correlation to this null distribution. D. Left panel, illustration of nucleotide importance scores for mesoderm and YSL CREs. The higher the letter, the greater its importance in predicting chromatin accessibility. The predicted TFBSs are highlighted in grey, and the motifs identified from ChIP-seq are displayed in the top right corner. Right panel: The ChIP-seq signal in either 2kb upstream or downstram of TFBS. The gray area indicates 95% confidence interval. E. Cumulative percentage of TFBSs from all motifs identified by TF-MoDISco within the top 10,000 cell-state-specific CREs. Note that over 80% of TFBSs come from the first three or four motifs. F. Distribution of the number of distinct motifs required to account for 80% of predicted TF binding sites for all 95 cell states. Note that for most of the 95 states, over 80% of TFBSs come from fewer than 6 motifs.

To extract sequence motifs that influence cell state accessibility predictions from DeepDanio, we used deep learning interpretation methods. Starting from the top 10,000 most specific putative CREs in each cell state, we used DeepExplainer (Lundberg & Lee, 2017) to calculate the contribution of each nucleotide to cell state predictions. We then used TF-MoDISco (Shrikumar et al., 2018) to identify, align, and cluster regions with high contribution scores into de novo motifs for each cell state. We identified an average of 17 motifs per cell state (see data access). Scanning the top 10,000 specifically accessible CREs for each cell state using TF-MoDISco-identified motifs provides cell-type-specific, genome-wide transcription factor binding site (TFBS) predictions. For example, nucleotides important for predicting accessibility in mesendoderm map to the motif for Tbxta (Figure 3D). For YSL, significant nucleotides for predicting accessibility emerge as a motif for Mxtx2 (Figure 3D). These results agree with previous findings that Tbxta and Mxtx2 are key players in the development of the mesoderm and YSL, respectively (Schier & Talbot, 2005; Schulte-Merker et al., 1994; Xu et al., 2012). We further validated the predicted TFBSs for the motifs that have available whole embryo ChIP-seq data (Nelson et al., 2017a; Xu et al., 2012), confirming that ChIP-seq signals are enriched at the predicted TFBS locations (Figure 3D and Figure S4B).

The above results indicate that DeepDanio can accurately predict chromatin accessibility and identify contributing motifs. To explore how many motifs are required to define cell-type-specific chromatin accessibility, we calculated the distribution of the predicted TFBSs for each motif across the top 10,000 specifically accessible CREs for each cell state. For many cell states, over 80% of TFBSs come from fewer than 6 motifs (Figure 3E-F). For example, among the YSL-specific CREs at 50% epiboly (onset of gastrulation), the Gata6 motif corresponds to 40% of the TFBSs, the Mxtx2 motif to another 30%, and the Hnf4 motif to another 10% (Figure 3E and Figure S4C). Many of these motifs are associated with TFs known to be fate regulators of the respective cell types. For example, Grhl3 is critical for EVL (Miles et al., 2017), Myod1 for adaxial cells (Weinberg et al., 1996), and Tp63 for the epidermis (H. Lee & Kimelman, 2002) (Figure 3E and Figure S4C). These results indicate that small sets of TFs play crucial roles in shaping the accessible chromatin landscape in each cell type (Wei et al., 2018).

### GRN reconstruction identifies critical regulators of cell identity

Since our data contain both gene expression and chromatin accessibility information from the same nuclei, it provides a way to incorporate CREs to infer enhancer-driven gene regulatory networks (eGRNs). We utilized the SCENIC+ pipeline (Bravo González-Blas et al., 2023) to reconstruct eGRNs during early zebrafish embryogenesis. Overall, SCENIC+ utilizes a three-step workflow (Figure 4A): First, we identified each TF’s potential CREs based on motif scanning using motifs collected in DANIO-CODE (Baranasic et al., 2022). Second, we linked these CREs to their target genes based on the correlation between gene expression and CRE accessibility. Third, we connected TFs to their target genes through CREs (see methods). This process resulted in a set of 100 enhancer-driven regulons (eRegulons), with each eRegulon consisting of a TF, its enhancers, and the target genes of these enhancers (Table S4). The 100 eRegulons contain 4,741 genes and 26,419 CREs. The size of each eRegulon varies from 10–976 genes, with a median size of 70 genes (Figure 4B).

**Figure 4:**
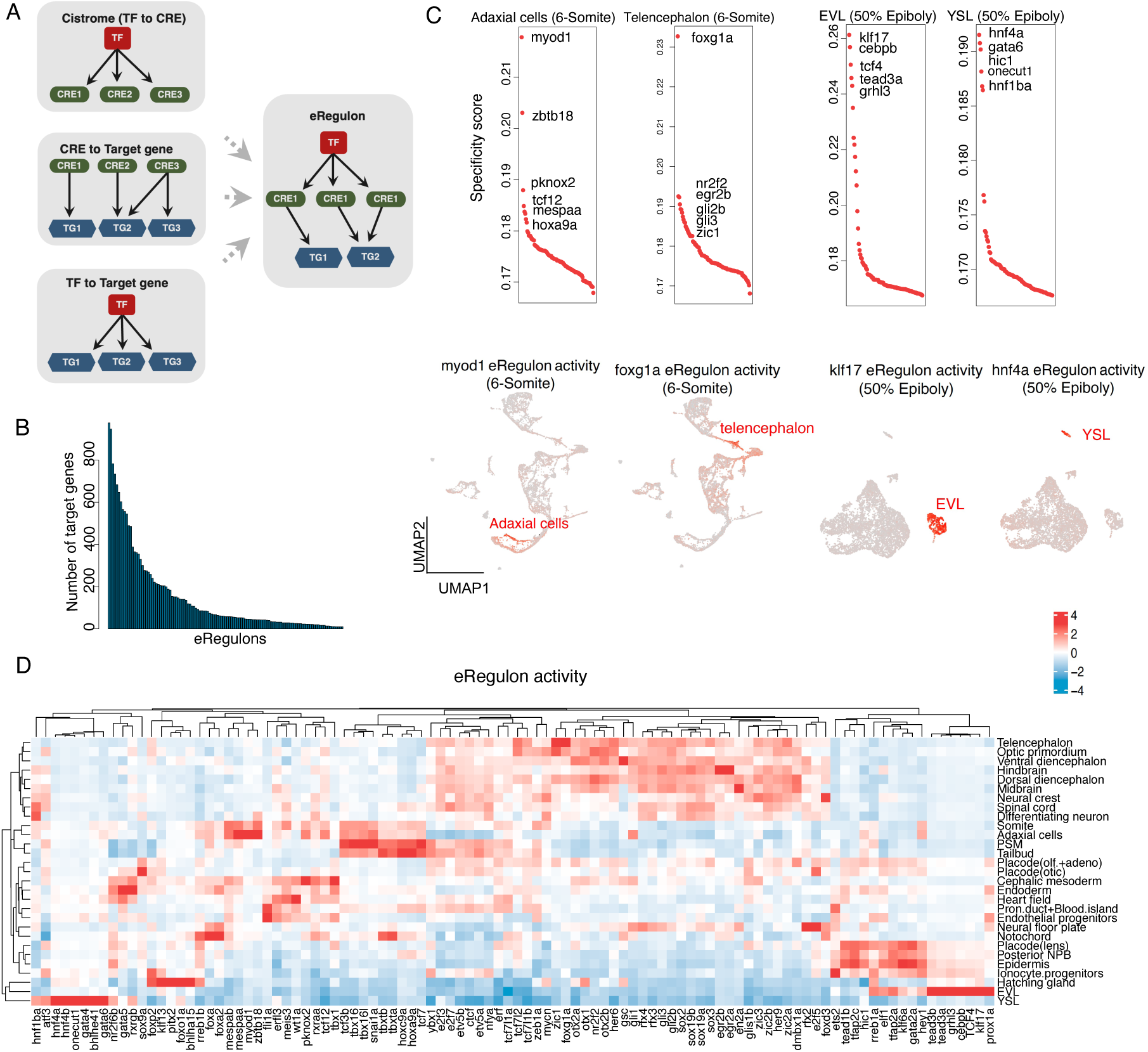
SCENIC+ infers enhancer-driven gene regulatory networks. A. SCENIC+ workflow. TF., transcription factor; CRE., *cis*-regulatory element; TG., target gene. A eRegulon consists of a TF, all the putative CREs regulated by this TF, and all the target genes regulated by this TF. B. Histogram of the number genes regulated by each TF. C. Upper panel: eRegulons ranked by specificity score. UMAP visualization showing eRegulon activity at the single-cell level, with cells colored according to their eRegulon activity levels. D. Heatmap displaying the mean eRegulon activity for all eRegulons across different cell types at the 6-somite stage. For each cell type, mean eRegulon activity was calculated by averaging the activity across all cells within the type. Hierarchical clustering was then performed based on these mean eRegulon activities to group cell types with similar activity patterns.

For each eRegulon, we evaluated its activities across 95 cell states (Table S5) using target gene expression and defined its cell state specificity score (Table S6) based on Jensen-Shannon divergence (see methods). The identified cell-state-specific eRegulons are consistent with current knowledge. For instance, the eRegulons specific to 6-somite adaxial cells include Myod1 (Weinberg et al., 1996) and Mespaa (Sawada et al., 2000), and the eRegulons specific to 6-somite telencephalon include Foxg1 (Zhao et al., 2009)and Nr2f2 (Chowdhury et al., 2024). At 50% epiboly (onset of gastrulation), the EVL-specific eRegulons include Klf17 (Liu et al., 2016; Miles et al., 2017) and Grhl3 (Miles et al., 2017), while the YSL-specific eRegulons include Hnf4a and Gata6 (Xu et al., 2012) (Figure 4C).

The UMAP plot provides additional support that the activities of these eRegulons are highly specific to their corresponding cell types (Figure 4C). Clustering cell types based on eRegulon activity shows that developmentally related cell types share similar eRegulon activity (Figure 4D). For instance, neural ectoderm cell types tend to cluster together, reflecting their shared eRegulon activity. Similarly, paraxial mesoderm cell types also exhibit clustering based on their shared regulatory activity. Taken together, our comprehensive network analysis connects accessible chromatin regions, putative enhancers, TFs and their binding motifs, and target genes.

### Combining GRNs and deep learning uncovers a shallow network driving instant differentiation

With the GRN, eRegulons and DeepDanio in hand, we wished to dissect the regulatory logic of the instant differentiation of EVL and YSL. We focused on the top 10 specific eRegulons in each stage of the EVL trajectory and YSL trajectory (Figure 5A). We discovered that the eRegulons associated with the EVL and YSL were remarkably large, including 12 of the 15 eRegulons with more than 360 target genes (Figure S5). For example, the two largest eRegulons, each including nearly 1000 genes, are Grhl3 in EVL and Onecut1 in YSL. We found that many of these eRegulons become active shortly after zygotic genome activation and reach peak activity at 50% epiboly (onset of gastrulation), in line with the observation that EVL and YSL have differentiated by this stage.

**Figure 5:**
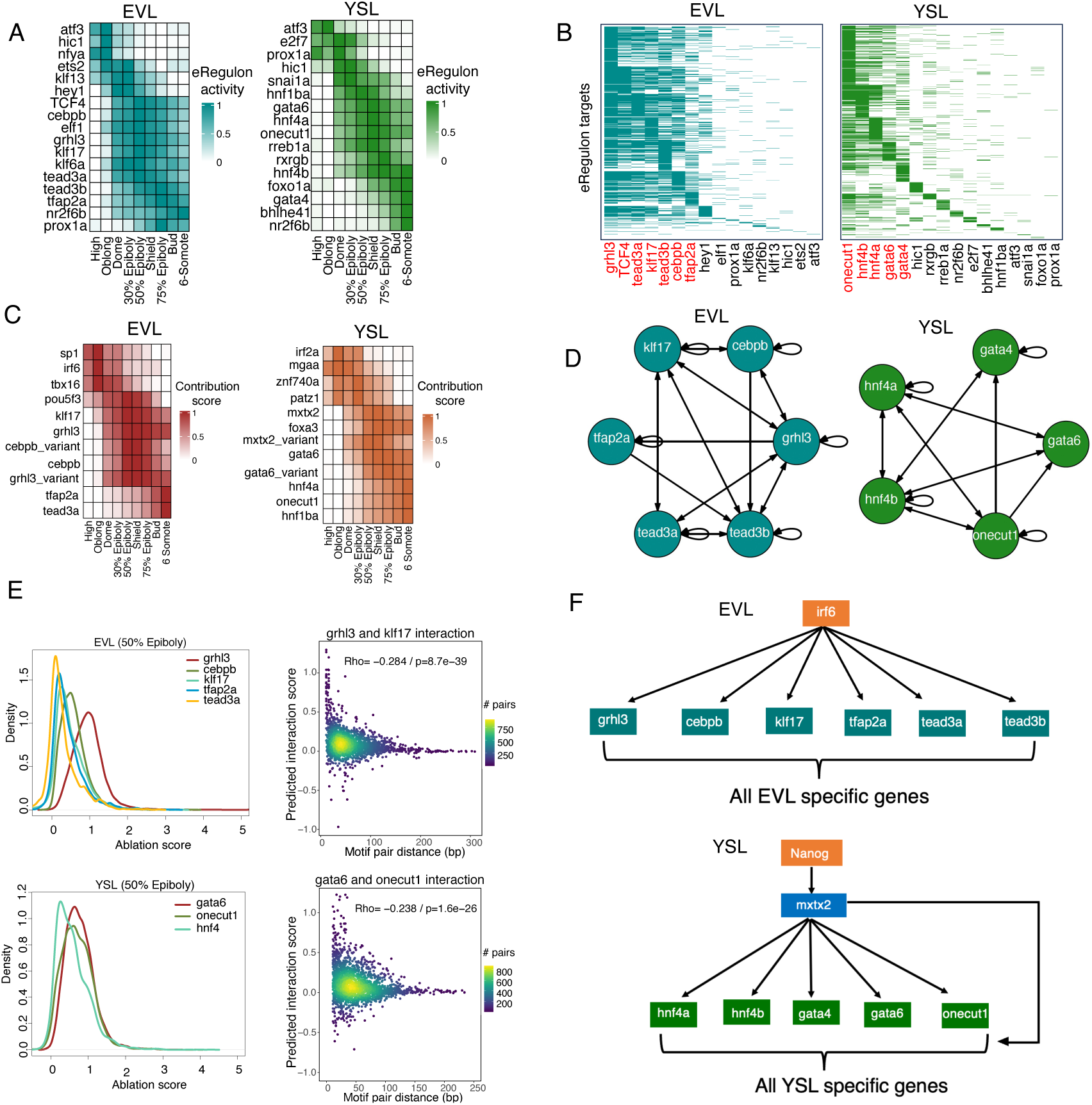
Gene regulatory logic underlying the instant differentiation of EVL and YSL. A. eRegulon activity dynamics for the EVL and YSL trajectories. The top 10 specific eRegulons in each cell state of the EVL (from oblong to 6-somite) and YSL (from dome to 6-somite) were selected, combined, and filtered to create a unique set of eRegulons. For each eRegulon, the mean activity across all cells within each cell state was then displayed. B. Overlap of target genes for all eRegulons displayed in panel A, showing that many targets are shared among a small set of TFs colored in red. C. TFBS contribution score dynamics for the EVL and YSL trajectories. The top 5 motifs identified by DeepDanio in each cell state of the EVL (from oblong to 6-somite) and YSL (from dome to 6-somite) were selected, combined, and filtered to create a unique set of motifs. For each motif, the mean contribution score across all its TFBSs was then displayed. D. Gene regulation among differentiation TFs, highlighting numerous auto-regulatory and cross-regulatory interactions. E. Motif ablation analysis. Left Panel: Distribution of ablation scores for each motif. Right Panel: Correlation between distances of two TFBSs and their interaction scores. The data show a trend where longer distances correspond to lower interaction scores. The Spearman’s correlation coefficient (Rho) and the P-value are displayed in the top-left corner of the plot. F. Topology of the gene regulatory network from maternally deposited TFs to differentiation genes. The initial input is maternally deposited TFs. Irf6 and Mxtx2 were not recovered using SCENIC+ because they are not included in the list of motifs in DANIO-CODE.

Plotting the targets of these eRegulons reveals that the majority of differentiation genes are regulated by only a few TFs (Figure 5B). Importantly, these TFs share many target genes, indicating that most differentiation genes are regulated combinatorially. By overlapping the TFs identified by SCENIC+ with the TFs identified by TF-MoDISCo and DeepDanio (Figure 5C and Figure S6), we defined high-confidence candidate TFs involved in the differentiation of EVL (Grhl3, Klf17, Cebpb, Tead3a, Tead3b, and Tfap2a) and YSL (Onecut1, Gata6, Gata4, Hnf4a, and Hnf4b). Notably, among these TFs, we also found auto-regulatory and cross-regulatory interactions. For example, Grhl3 can regulate itself and also engage in cross-regulation with Klf17, Cebpb, Tead3a and Tead3b (Figure 5D). Overall, the inferred GRN suggests that instant differentiation is regulated by a handful of TFs that function together to activate a large number of differentiation genes.

To explore this potential combinatorial regulation further, we conducted in-silico ablation(Yin et al., 2024) analysis of the motifs from DeepDanio. In this analysis, motifs were replaced with dinucleotide shuffles to preserve sequence composition while disrupting the motif. The resulting decrease in accessibility was quantified as an ‘ablation score’ (see methods), with higher scores indicating a greater impact on chromatin accessibility. Our findings reveal that while all motifs influence chromatin accessibility, Grhl3 has a much higher impact than the others, aligning with previous research that categorizes it as a pioneer TF(Jacobs et al., 2018)(Figure 5E). To measure the interaction score for pairs of motifs co-occurring on the same CRE, we calculated the sum of two individual motif ablation scores minus the score from ablating both motifs simultaneously. A score above zero indicates cooperativity between co-binding TFs, contributing to chromatin accessibility. For example, we discovered that the majority of motif pairs between Grhl3 and Klf17 exhibit scores greater than zero (Figure 5E), supporting the existence of cooperative regulation between the two TFs. Additionally, we observed that this regulatory cooperation is influenced by the distance between motifs, with the interaction score decreasing as the distance increases (Figure 5E). Other motif pairs show similar patterns as Grhl3 and Klf17 (Figure S7). These simulations suggest that there is extensive distance-dependent cooperativity between co-binding TFs.

To identify the putative regulators that might activate these EVL and YSL differentiation TFs, we first identified DeepDanio motifs exhibiting high contribution scores at the onset of differentiation (Figure 5C). We then narrowed our focus to motifs whose corresponding TFs are expressed in the EVL or the YSL. Following these criteria, we identified two primary candidates: Irf6 for EVL and Mxtx2 for YSL (Figure 5F). This finding aligns with the important roles of these factors in the EVL and YSL and suggests that Irf6 and Mxtx2 activate the differentiation TFs to initiate EVL and YSL differentiation (Liu et al., 2016; Sabel et al., 2009; Xu et al., 2012). Further inspection of putative Mxtx2 TFBSs showed that these motifs are not only found in differentiation TF genes but also in their target genes. Moreover, Mxtx2 TFBSs frequently co-localize with the TFBSs of the differentiation TFs (gata6, hnf4, and onecut1; Figure S8). This prediction suggests that Mxtx2 functions in a feedforward loop in which it activates differentiation TFs and then together with differentiation TFs activates their target genes (Figure 5F). Notably, the expression of *mxtx2* is driven by Nanog (Xu et al., 2012), a maternally deposited TF, as is Irf6 (Sabel et al., 2009). These observations indicate that the GRN from pluripotency to the terminal differentiation of EVL and YSL is very shallow (Figure 5F).

### Mis-expression of differentiation transcription factors inactivates hundreds of target genes

The SCENIC+ and DeepDanio analyses predict that a small set of differentiation TFs (Grhl3, Klf17, Cebpb, Tead3a, Tead3b, and Tfap2a) activates a large number of EVL differentiation genes (Figure 5F). To test this prediction experimentally, we mis-expressed these TFs and performed scRNA-seq at 50% epiboly (onset of gastrulation, when the EVL has formed, Table S8). Strikingly, the differentiation TFs ectopically activated hundreds of EVL differentiation genes. Compared to the mCherry control (Figure 6A), we observed the emergence of a new gene expression cluster that expresses EVL genes. For example, *cldne* is expressed in both the EVL and the “EVL-like” clusters upon mis-expression of differentiation TF (Figure 6A). We found that almost all SCENIC+ predicted EVL-specific target genes are highly expressed in the EVL-like cells (Figure 6B). In addition, EVL-like cells and deep cells (mesendodermal and ectodermal cells) had distinct gene expression profiles, indicating that EVL-like cells are not in a hybrid state but transcriptionally resemble EVL cells (Figure 6B). Overall, these results demonstrate that the differentiation TFs are sufficient to activate EVL differentiation genes.

**Figure 6:**
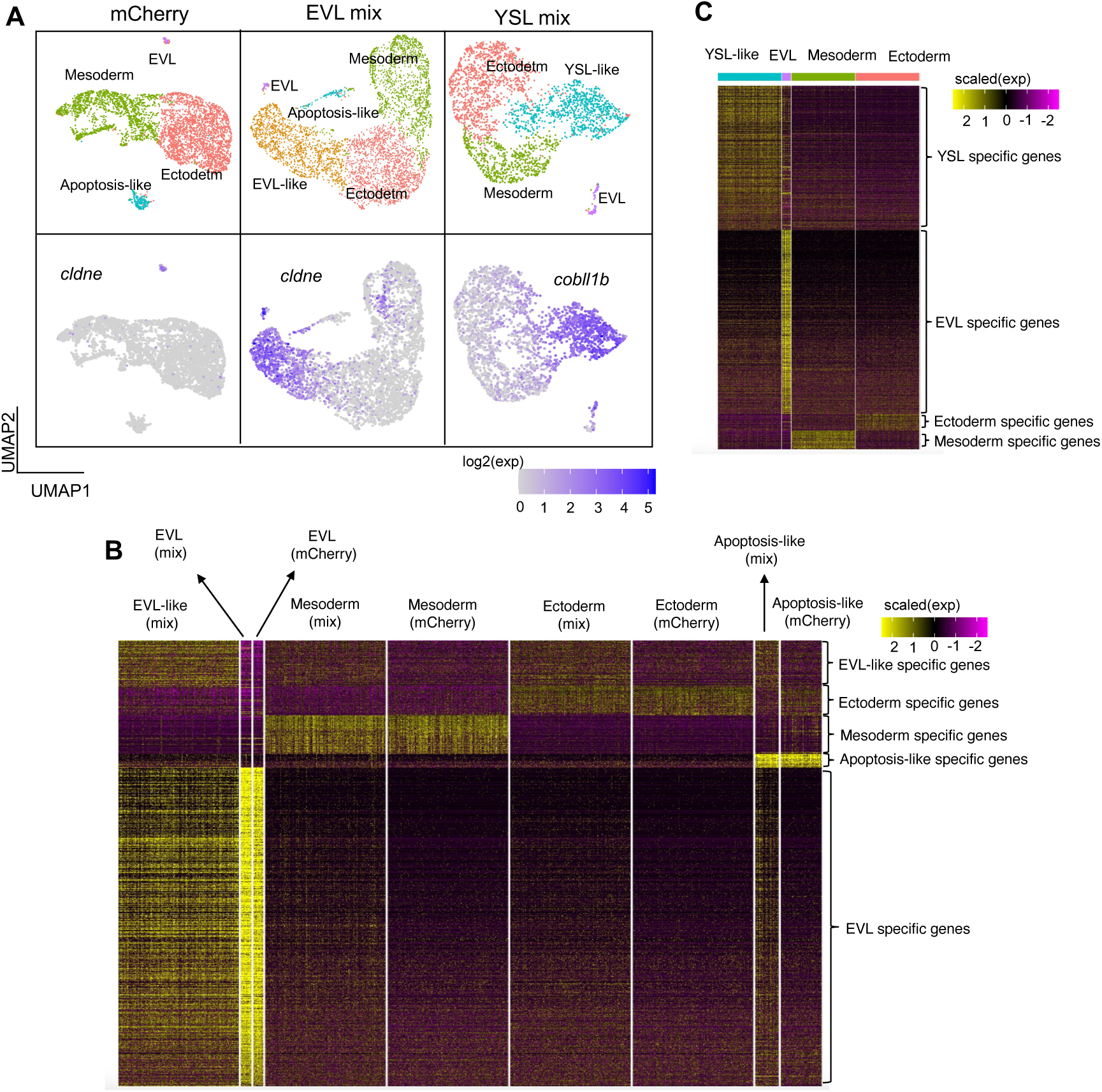
Mis-Expression of differentiation transcription factors ectopically activates EVL and YSL differentiation genes. A. UMAP projection of single-cell transcriptomes, with cells colored by cell type (upper panel) and by marker gene expression (lower panel). *cldne* and *cobll1b* are used as markers for EVL and YSL, respectively. In addition to the native EVL, we identified an “EVL-like” cluster expressing EVL markers. It should be noted that since we performed single-cell RNA-seq rather than single-nucleus RNA-seq, we couldn’t capture YSL cells as they are multinucleated and exist in a syncytium However, a “YSL-like” cluster was identified, expressing YSL marker genes. B. Heatmap showing gene expression of cell type-specific genes across different cell types under two conditions: EVL mix (mis-expression of all differentiation TFs) and EVL mCherry (mis-expression of mCherry). Notably, most EVL-specific genes are activated in the “EVL-like” cluster, although their expression levels are lower than in native EVL. The 77 “EVL-like” specific genes refer to those highly expressed in the “EVL-like” cluster (compared to native EVL from the mCherry control, fold change (log2) > 1, p-value < 0.01). These genes are typically weakly expressed housekeeping genes during normal development and are primarily involved in RNA biosynthetic processes. C. Heatmap showing gene expression of cell type-specific genes across different cell types under the YSL mix condition (mis-expression of all differentiation TFs plus Mxtx2). Most YSL-specific genes are activated in the “YSL-like” cluster.

We performed the same experiment for the YSL and mis-expressed Mxtx2 together with the differentiation TFs (Onecut1, Gata6, Gata4, Hnf4a, and Hnf4b). TF mis-expression ectopically activated almost all SCENIC+ predicted YSL differentiation genes (Figure 6C). To further test the feedforward role of Mxtx2 predicted by DeepDanio, we mis-expressed the differentiation TFs without Mxtx2 and found that the majority of YSL-specific genes were not activated in deep cells, supporting a feedforward loop involving Mxtx2 (Figure S9).

In summary, the mis-expression of differentiation TFs identified by SCENIC+ and DeepDanio ectopically activates hundreds of EVL and YSL differentiation genes. The transformed cells do not express genes specific to other cell types and express the large majority of EVL- or YSL-specific genes. These results indicate that a shallow network of differentiation TFs underlies the process of instant differentiation.

## Discussion

In this study, we generated and explored a high-quality single-cell multi-omics atlas for early zebrafish embryogenesis (Figure 1 and Table S1). The atlas provides a rich resource for future studies of zebrafish development, including integration with spatial transcriptomics (Wan et al., 2024) and in-toto live imaging (McDole et al., 2018). The reconstructed GRN and eRegulons will help dissect the regulatory logic of cell type specification and differentiation. In addition, the DeepDanio deep learning model developed here can be applied to dissect cell-type- or stage-specific enhancer architecture and design synthetic enhancers with specific temporal and spatial activities (de Almeida et al., 2024; Taskiran et al., 2024).

Our study also presents a generalizable framework to dissect the regulatory logic of embryonic specification and differentiation through integrating single-cell multi-omics, deep learning, GRN reconstruction and in vivo assays. We provide three main insights:

First, we defined the emergence of accessible chromatin regions and their relationships with gene expression. We identified three distinct temporal waves of dCRE expansion: EVL and YSL > mesendoderm > neuroectoderm (Figure 1D). This pattern closely follows the emergence of these cell types but contrasts with the early accessibility of promoters (Figure 1E), which mark genes for later expression (Pálfy et al., 2020; Reddington et al., 2020). Comparing the temporal emergence of dCRE with the timing of gene activation revealed that dCRE priming (chromatin is accessible before gene transcription) is common: mesendodermal dCREs are primed in pluripotent cells at the high stage, while neuroectodermal dCREs are primed at the 50% epiboly stage (onset of gastrulation) (Figure 2B and 2C). This early priming of mesendodermal dCREs might facilitate mesendoderm specification during gastrulation by enabling a rapid response to inductive signals such as Nodal (Schier & Talbot, 2005). Notably, we discovered that hundreds of mesendodermal dCREs were primed by Nanog (Figure 2E and 2F). This pioneer TF is known to activate several hundred genes to promote pluripotency (Boyer et al., 2005; M. T. Lee et al., 2013). Our results reveal that Nanog has a dual role: it promotes pluripotency while also prepares the embryo for subsequent mesoderm induction. These results extend and generalize studies of individual genes found to be primed in flies (Falo-Sanjuan et al., 2019), frogs (Charney et al., 2017), and human and mouse embryonic stem cells (Kim et al., 2018; Liber et al., 2010; A. Wang et al., 2015). Our results contradict an earlier report that suggested that mesendodermal dCREs are not primed in the mouse epiblast (Argelaguet et al., 2019), but align with a recent preprint that does suggest priming (Sendra et al., 2024).

Second, the predictions from DeepDanio suggest that a small number of TFs play a critical role in shaping the accessible chromatin landscape (Figure 3). This observation might seem surprising considering the broad array of TFs typically expressed in cells, but it complements a recent study (Wei et al., 2018) that developed a massively parallel protein activity assay to measure the DNA-binding activity of all TFs in cell or tissue extracts through electrophoretic mobility shift assays (EMSAs). Only a small set of TFs displayed strong DNA-binding activity, and the sequence features underlying the DNA-binding sites of these TFs can accurately predict the cell type-specific accessible chromatin landscape. Collectively, these results support the idea that a limited number of TFs establish the overall chromatin landscape and thus facilitate the binding of TFs with weaker chromatin-modulating capabilities.

Third, we discovered a shallow gene regulatory network underlying instant differentiation: an upstream TF activates a set of differentiation TFs, which in turn activate hundreds of effector genes that define cell-type-specific structures and functions. In the case of the EVL, the maternally deposited TF Irf6 serves as the upstream TF that directly activate multiple differentiation TFs to initiate differentiation (Figure 5F). In the case of the YSL, Nanog is the upstream TF that activates Mxtx2, which then activates differentiation TFs. Together, Mxtx2 and these differentiation TFs form a feedforward loop to initiate differentiation. Since the expression of *mxtx2* is transient (Figure S10), its feedforward role may involve cooperation with the differentiation TFs to open the chromatin of enhancers in the YSL-specific genes (Figure S11). The differentiation TFs then activate and maintain the expression of differentiation genes.

This shallow network model suggests that instant differentiation occurs because differentiation TFs are quickly activated by maternally deposited TFs, thereby skipping multiple cell fate transitions typically found during the stepwise differentiation of cell (Davidson, 2010; Liberali & Schier, 2024; Murtaugh, 2007; Y. Wang et al., 2023). Indeed, TF mis-expression analysis showed that the identified EVL and YSL differentiation TFs are sufficient to ectopically activate nearly all effector genes at 50% epiboly (onset of gastrulation, when the EVL and YSL have formed). These findings support the proposed shallow network model and extend previous studies (De La Garza et al., 2013; Liu et al., 2016; Sabel et al., 2009; Xu et al., 2012) that examined the effects of some differentiation TFs on a few marker genes.

While direct activation of effector genes by maternal TFs could theoretically speed up differentiation, the EVL/YSL regulatory design offers advantages. Activating multiple differentiation TFs enhances the specificity of effector gene expression through cooperative regulation, as indicated by in-silico motif ablation analysis (Figure 5E). Additionally, we observed auto-regulatory and cross-regulatory interactions among these TFs, similar to those in embryonic stem cell networks that help maintain stemness (Boyer et al., 2005) (Figure 5D). Given the transient expression of maternal factors, these interactions might sustain and amplify effector gene expression, ensuring sufficient production of the gene products necessary for differentiation.

The observation that cell types like the EVL and YSL can undergo instant differentiation suggests that stepwise differentiation programs might not be necessary for differentiation TFs to activate their effector genes. Future studies will address the necessity of stepwise differentiation: Can stepwise differentiation be transformed into instant differentiation?

## Methods

### Culturing and collecting embryos

Embryos from wild-type (Tupfel longfin/AB) crosses were collected 20 minutes after fertilization. They were dechorionated by incubation in 1 mg/ml pronase (Sigma-Aldrich) for 5 minutes until chorions began to blister, submerged in ∼200ml of zebrafish “blue water” embryo medium (5 mM NaCl, 0.17 mM KCl, 0.33 mM CaCl2, 0.33 mM MgSO4, 0.1% methylene blue, dissolved in fish system water) in a glass beaker, and then the blue water was decanted and vigorously replaced 3 times. It was critical to prevent embryos from contacting air or plastic during this process or at any point thereafter. Embryos were then cultured at 28°C in blue water in plastic Petri dishes that had previously been coated with 2% agarose (dissolved in blue water).

### Cell dissociation and nuclei isolation

Single cell suspensions were obtained using an embryo fractionation protocol. Approximately 200 dechorionated embryos for each experimental condition were transferred into a 2 ml LoBind Eppendorf tube (EP-022431048) containing 1.5 ml of chilled deyolk buffer (55 mM NaCl, 1.8 mM KCl, 1.25 mM NaHCO3). The embryos were gently pipetted up and down 3-4 times using a p1000 tip and then placed in a thermal shaker at 1200 rpm at 4°C for 20 seconds. The tubes were centrifuged at 4°C at 250g for 4 minutes. After centrifugation, the supernatant was discarded, and 1 ml of deyolking wash buffer (10 mM Tris pH 8.5, 110 mM NaCl, 3.5 mM KCl, 2.7 mM CaCl2) was added to each tube to resuspend the pelleted embryos. Resuspension was performed by pipetting six times with a p1000 tip. The tubes were then centrifuged again, the supernatant discarded, and the cell pellets resuspended in 1 ml DMEM (Gibco, 11594426) with 0.1% BSA. This centrifugation and resuspension step with DMEM (0.1% BSA) was repeated once more. After another round of centrifugation, 150 μl of chilled 0.5X Lysis Buffer (10 mM Tris pH 7.5, 10 mM NaCl, 3 mM MgCl2, 1% BSA, 0.1% Tween, 1 mM DTT, 1 U/μl RNaseIn (Promega), 0.1% NP40, 0.01% Digitonin) was added to the pellet. The mixture was pipetted 5 times with a p200 tip and incubated on ice for 5 minutes. Next, 1.5 ml of chilled Wash Buffer (identical to NE buffer but without NP40 and digitonin) was added to the lysed cells. The cells were pipetted five times and then centrifuged at 450g for 5 minutes at 4°C. The supernatant was carefully removed without disturbing the nuclei pellet. The pellet was resuspended in at least 300 μl of chilled Diluted Nuclei Buffer (10x Genomics). The suspension was then passed through a Flowmi Cell Strainer (40 μm). Finally, the nuclei concentration and morphology were checked using a hemocytometer.

### 10x Multiome library preparation and sequencing

Nuclei were diluted to ensure a maximum of 8,000 were used for the 10x Multiome library preparation. For the high stage, oblong, dome, and 30% epiboly stages, we have one technical replicate each. For the 50% epiboly and 6-somite stages, we have two technical replicates each. For the shield, 75% epiboly, and bud stages, we have two biological replicates, and for each biological replicate, we have two technical replicates. Libraries were prepared from the single nuclei suspensions using the 10x Chromium Next GEM Single Cell Multiome ATAC + Gene Expression kit, following the standard 10x protocol. The libraries were sequenced on a NovaSeq platform (Illumina) using the recommended read lengths. This sequencing yielded an average of 348 million RNA-seq reads and 536 million ATAC reads per sample. We recovered an average of 2,500 cells per sample prior to quality control.

### scRNA data and scATAC data processing

Raw sequencing files were processed with CellRanger arc 2.0.0 using default parameters. Reads were mapped to the Ensembl GRCz11 reference genome (Ensembl Release 100)(Yates et al., 2020). Low-quality cells were filtered out based on several quality control metrics: the number of expressed genes, the proportion of mitochondrial reads for RNA; the number of fragments, and the TSS enrichment score for ATAC. For RNA: For the high and oblong stages, since they mark the onset of transcription, the number of expressed genes is lower than in later stages. Therefore, we filtered out cells with fewer than 200 (low quality cells) or more than 3000 (to exclude potential doublets) expressed genes. For later stages, we filtered out cells with fewer than 500 or more than 5000 expressed genes. We also excluded cells with more than 10% of reads mapping to the mitochondrial genome. For ATAC: We filtered out cells with a TSS enrichment score lower than 4. We also excluded cells with fewer than 1000 unique nuclear fragments.

### Cell clustering and annotation

The scRNA-seq part of the scMultiome data was processed and analyzed using Seurat version 4.1.0(Hao et al., 2021). We processed each developmental stage separately and clustered the cell types accordingly. Marker genes for each cell cluster were identified using the Wilcoxon test, comparing the expression levels within the cluster to those in the rest of the cells. To annotate the cell types represented by each cluster, we checked the expression profiles of the significant marker genes with high fold changes against the ZFIN database and our previously constructed developmental trajectories.

### Peak calling

The scATAC-seq part of the scMultiome data was processed and analyzed using ArchR version 1.0.1(Granja et al., 2021). Using the cell types identified from the scRNA-seq data as groups, pseudo-bulk replicates were generated for each cell type. Peak calling was then performed using MACS2 version 2.2.7.1 through ArchR with the following parameters: “--shift -75 --extsize 150 --nomodel --call-summits --nolambda --keep-dup all -q 0.01”. We did not call peaks for cell types with fewer than 80 cells. These excluded cell types were: YSL(dome), YSL(30% epiboly), YSL(6-somite), forerunner cells(75% epiboly), ionocyte progenitors(6-somite), endothelial progenitors(6-somite), neural floorplate(6-somite). Calling peaks in a cell type-aware manner resulted in multiple peak sets that needed to be consolidated into a single peak annotation. Using ArchR’s iterative overlap merging procedure, we obtained a consensus peak set consisting of 444,653 peaks, each 500 bp wide. Distal CREs (dCREs), including putative enhancers, were defined as peaks >500 bp from an annotated transcription start site (TSS). Peaks within <500 bp of TSS were annotated as promoters.

### 10x standalone scATAC-seq library preparation and sequencing

To compare scATAC-seq data from scMultiome with standalone scATAC-seq, we also generated standalone scATAC-seq data for two stages: 50% epiboly and the 6-somite stage. For the 50% epiboly stage, we have two technical replicates, and for the 6-somite stage, we have three technical replicates. Libraries were prepared from the single nuclei suspensions using the Chromium Next GEM Single Cell ATAC Reagent Kits v1.1, following the standard 10x protocol. The libraries were sequenced on a NovaSeq platform (Illumina) using the recommended read lengths. This sequencing yielded an average of 368 million ATAC reads per sample. We recovered an average of 4’010 cells per sample prior to quality control.

### Cellular trajectories reconstruction

To connect each cell cluster observed at a given stage with its pseudoancestor in the preceding stage, we applied a k-NN (k-nearest neighbors) method as described in earlier studies. First, we merged all cells from the given stage and the preceding stage using Seurat (version 4.1.0) and projected them into a common UMAP embedding space. We then calculated the Euclidean distances between individual cells from the given stage and the preceding stage within this UMAP space. For each cell cluster at the given stage, we identified their five nearest neighbors from the preceding stage and calculated the proportion of these neighbors derived from each cell cluster in the preceding stage. This process was repeated 500 times with 80% subsampling from the same embedding. The median proportions of neighbors were then used as weights for edges between a cell cluster and its potential antecedents. Only edge weights greater than 0.2 were retained (Table S2) for constructing the resulting acyclic directed graph, which is shown in Figure 1B.

### CRE usage during embryogenesis

We applied the AccessiblePeaks function from the Signac package to identify accessible CREs in different cell types. A CRE is considered “accessible” in a cell type if it is accessible in at least 5% of the cells of that cell type(Kelley et al., 2016; Sarropoulos et al., 2021; Stuart et al., 2021). For each trajectory analysis (Figure 1C and D), at each stage, we examined how many CREs became accessible and whether their accessibility was maintained up to the 6-somite stage.

### The timing differences between CRE chromatin opening and target gene activation

#### 1. Identify cell type specific genes

For cell types at the 50% epiboly stage (EVL, YSL, dorsal anterior, dorsal posterior, ventrolateral mesendoderm, and margin tail), we identified the genes specifically expressed in each cell type using the Seurat FindMarkers function (fold change (log2) > 1, p-value < 0.01). We then filtered out genes that were already expressed at the oblong stage. A gene is considered expressed in a cell type if it is expressed in at least 10% of the cells in that cell type(Bravo González-Blas et al., 2023; Karlsson et al., 2021; Kotliar et al., 2019; Meng et al., 2024). Similarly, for cell types at the bud stage (forebrain, midbrain, hindbrain, epidermis, hatching gland, notochord, tailbud, lateral plate mesoderm, presomitic mesoderm, adaxial cells, and endoderm), we identified the genes specifically expressed in each cell type using the Seurat FindMarkers function (fold change (log2) > 1, p-value < 0.01). We then filtered out genes that were already expressed at the shield stage.

#### 2. Infer the putative distal CREs (dCREs) of cell type specific genes

For each cell type at the 50% epiboly stage, we extracted the cells along the trajectory from the high stage to the 50% epiboly stage. For each cell type at the bud stage, we extracted the cells along the trajectory from the 50% epiboly stage to the bud stage. We then used the LinkPeaks function from Signac to infer the putative dCREs of each gene. The LinkPeaks function computes the Pearson correlation coefficient (r) between gene expression and chromatin accessibility for each dCREs located within 50 kb upstream or downstream of the transcription start site (TSS). This analysis includes all cells extracted from the trajectory. For each dCREs, LinkPeaks also computes a background set of expected correlation coefficients by randomly sampling 200 CREs located on a different chromosome from the gene, matched for GC content, accessibility, and sequence length to estimate a p-value. An associated dCRE of a gene is defined as having a Pearson correlation coefficient > 0.1 and a p-value < 0.05.

#### 3. Calculate the proportion of accessible CREs in each stage

For the cell type-specific genes in each cell type, we calculated the proportion of their promoters or dCREs that are accessible at each stage of the trajectory as shown in Figure 1E.

### Enhancer priming analysis

#### 1. Infer the associated distal CREs (dCREs) for mesendodermal and neuroectodermal genes

For mesendodermal cell types at the 50% epiboly stage (dorsal anterior, dorsal posterior, ventrolateral mesendoderm, and margin tail), we identified genes specifically expressed in each cell type using the Seurat FindMarkers function (fold change (log2) > 1, p-value < 0.01). We then filtered out genes already expressed at the oblong stage. A gene is considered expressed in a cell type if it is expressed in at least 10% of the cells in that cell type(Bravo González-Blas et al., 2023; Karlsson et al., 2021; Kotliar et al., 2019; Meng et al., 2024). Similarly, for neuroectodermal cells at the bud stage (forebrain, midbrain, hindbrain), we identified genes specifically expressed in each cell type using the Seurat FindMarkers function (fold change (log2) > 1, p-value < 0.01). Genes already expressed at the shield stage were filtered out. For each gene, we identified dCREs located within 50 kb upstream or downstream of the transcription start site (TSS). A dCRE was considered associated with a gene if the Pearson correlation coefficient (association score) between gene expression and dCRE chromatin accessibility was > 0.1, with a p-value < 0.05. For more details, refer to “**Infer the putative distal CREs (dCREs) of cell type specific genes**”.

#### 2. Compare the proportion of primed dCREs between associated and non-associated dCREs, as shown in Figure 2B and C

A primed dCRE at a given stage is defined as a dCRE that is accessible at that stage, but its target gene is not yet expressed. We applied the AccessiblePeaks function from the Signac package to identify accessible CREs in a given cell type. Check “CRE usage during embryogenesis” for details. A gene is considered expressed in a cell type if it is expressed in at least 10% of the cells in that cell type(Bravo González-Blas et al., 2023; Karlsson et al., 2021; Kotliar et al., 2019; Meng et al., 2024).

#### 3. Chromatin accessibility analysis

The chromatin accessibility data (bedGraph file) comes from Miao et al., (2022). They performed bulk ATAC-seq at the sphere stage (4 hpf) in nanog, pou5f3, and sox19b triple mutants (MZnps) condition, wild type condition, as well as in various rescue conditions by injecting mRNAs into MZnps embryos. For each primed and non-primed dCREs, we calculated the chromatin accessibility in different conditions using the bedtools “map” command(Quinlan & Hall, 2010), as shown in Figure 2E.

#### 4. Nanog binding site analysis

The Nanog binding peaks, identified using CUT&RUN, come from Wang et al., (2022). For each dCRE, if it overlapped with a Nanog binding peak by at least 1 bp, we defined this dCRE as binding with Nanog. Then, we calculated the proportion of primed and non-primed dCREs with Nanog binding, as shown in Figure 2F.

### Deep learning analysis

#### 1. Data preprocessing

We started from a 444,653 x 95 matrix containing aggregated Counts Per Million (CPM) for each scATAC peak and cell state. Log10(CPM) values were then calculated with a per cell-state pseudocount obtained from the corresponding minimum non-zero CPM value. Finally, quantile normalization was applied across cell states. 500nt-long peak sequences were extracted from the GRCz11 version of the Danio Rerio genome. We additionally extracted 200,000 500nt-long fragments from random non-peak genome locations (negative samples).

#### 2. Training splits

We distributed all 25 Danio Rerio chromosomes into 10 subsets with a roughly similar number of scATAC peaks (mean: 44.5 thousand peaks per subset) using the prtpy python package (https://github.com/coin-or/prtpy). For each individual model in the DeepDanio ensemble, eight subsets were used directly for model fitting (training), one was used for early stopping (validation), and one was held out from training for performance evaluation (test). Each model was assigned a different test set. The list of chromosomes used in each model can be found in Table S7.

#### 3. Training

DeepDanio is an ensemble of three residual neural networks as shown in Figure 3A. The input is a one hot-encoded 500bp sequence. Outputs are predicted log10CPM for each of the 95 cell states. Model training was performed with python 3.10 and tensorflow 2.10 using AWS EC2 g5.2xlarge instances. The loss was the sum of a standard MSE component and a per-sequence Pearson r component that incentivizes learning the relative order of predicted log10 CPMs across cell states. More specifically, we adapted the AI-TAC Pearson r loss implementation(Maslova et al., 2020)where we maximize the cosine similarity between the per-sequence mean-normalized prediction and the mean-normalized measurement. Before training, positive (i.e. the preprocessed scATAC matrix) and negative data were split into training/validation/test sets according to chromosomes as described above. Training data was further augmented with the reverse complement of every sequence. At the beginning of each epoch, we randomly selected enough negative samples to reach a 5:1 ratio of positive-to-negative training peaks. Negative peaks were assigned the lowest log10CPM value on each cell state. We used the Adam optimizer with default settings, a learning rate of 2e-4, and early stopping on the validation loss with a patience of two. We included negative samples for validation loss evaluation but no reverse complements. Once training finished, we performed another training round starting from the optimized weights, which resulted in a slight additional performance improvement. Three models were independently trained on different chromosome splits as described above. For performance evaluation (i.e. Figure 3B and C and Figure S4A) we aggregate predictions of each model over its own test set. For all other analyses in this manuscript, we used the mean of all three models.

#### 4. De novo motif discovery

We extracted motifs that contribute to accessibility in each cell state using the contribution scores of the 10,000 most cell state-specific peaks. We first filtered out peaks where the square prediction error averaged across cell states was 0.125 or greater, retaining 377,312 peaks (84.86%). Then, for each cell state, we obtained the most specific peaks by sorting by their predicted specificity score, defined as the predicted log10CPM in the target cell state minus the average predictions across all other states, and retained the top 10,000. Next, we calculated nucleotide contribution scores of each set of cell state-specific peaks with respect to the model output corresponding to that cell state. We used a custom version of DeepExplainer/DeepSHAP (https://github.com/castillohair/shap/tree/castillohair/genomics_mod) modified to work with tensorflow 2 and generate hypothetical contribution scores needed for TFModisco. As background, we used 10 random dinucleotide shufflings per sequence. Next, we ran TFModisco (Shrikumar et al., 2018), specifically the modisco-lite implementation (https://github.com/jmschrei/tfmodisco-lite), on the contributions of each set of cell state-specific peaks, with 25,000 as the maximum number of seqlets per metacluster. This resulted in an average of 17 motifs per cell state (Table S3), each with a corresponding PWM and contribution weight matrix (CWM). Finally, to generate the plots in Figures 3-F, we realigned motifs to their corresponding cell state-specific peaks using the motif scan method in Avsec et al (Avsec et al., 2021)which considers nucleotide contributions in addition to motif PWM similarity.

#### 5. Validation of deep learning-identified transcription factor binding sites (TFBS) using ChIP-Seq data

The ChIP-seq data for nanog performed at the high stage were obtained from Xu et al.(Xu et al., 2012). In this study, C-terminally myc-tagged zebrafish nanog was overexpressed, and myc antibodies were used for immunoprecipitation; The ChIP-seq data for pou5f3 performed at the sphere stage were obtained from Miao et al. ((Miao et al., 2022). Similarly, C-terminally myc-tagged zebrafish pou5f3 was overexpressed, and myc antibodies were used for immunoprecipitation. The ChIP-seq data for mxtx2 performed at the dome stage were obtained from Xu et al. (Xu et al., 2012). Here again, C-terminally myc-tagged zebrafish mxtx2 was overexpressed, and myc antibodies were used for immunoprecipitation. The ChIP-seq data for tbxta performed at the 75% epiboly stage were obtained from Nelson et al. (Nelson et al., 2017). In this case, anti-tbxta antibodies were used for immunoprecipitation. The ChIP-seq data for cdx4 performed at the bud stage were obtained from Paik et al. (Paik et al., 2013). C-terminally myc-tagged zebrafish cdx4 was overexpressed, and myc antibodies were used for immunoprecipitation.

For nanog and pou5f3, the deep learning-identified TFBS were derived from the high stage. For mxtx2, the TFBS were identified in 50% epiboly YSL cells. For tbxta, the TFBS were found in 50% ventrolateral mesoendoderm cells. For cdx4, the TFBS were identified in the lateral plate mesoderm, presomitic mesoderm, and tailbud cells at the bud stage, as cdx4 is expressed in all three cell types. For each TFBS identified through deep learning, the region was extended by 2 kb on either side. These extended regions were then divided into 20 bp bins, and the ChIP-seq signal (bedGraph file) for each bin was calculated using the bedtools “map” command(Quinlan & Hall, 2010), as shown in Figure 3D and Figure S4B.

#### 6. Motif in-silico ablation analysis

For each TF involved in EVL differentiation (grhl3, cepbp, klf17, tfap2a, and tead3a) and YSL differentiation (mxtx2, gata6, and hnf4), we first performed single TFBS ablation analysis, as shown in Figure 5E left panel and Figure S11A. If the TFBSs of one TF colocalized with the TFBSs of other TFs within the same dCREs, we also conducted double ablation analysis for each pair of co-binding TFBSs. For ablation analysis, the identified TFBSs were replaced with dinucleotide shuffles to preserve sequence composition while disrupting the motif. Chromatin accessibility of both the original and ablated sequences was calculated using DeepDanio. Each ablation was repeated 10 times, and the mean chromatin accessibility value was computed for the ablated sequences. The ablation score was calculated using the formula:

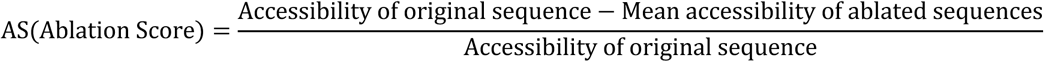

To study pairwise motif interactions, as shown in Figure 5E (right panel), Figure S7, and Figures S11B, C, D, we calculated ablation scores for single motif ablations (AS(A) for motif A and AS(B) for motif B) and for double ablations (AS(AB)). We then compared the sum of the individual ablation scores with the double ablation score to determine the type of interaction:

Additive Effect: AS(A)+ AS (B) = AS (AB) Redundant Effect: AS (A)+ AS (B) < AS (AB) Cooperative Effect: AS (A)+ AS (B) > AS (AB)

### GRN analysis

We followed the tutorial from SCENIC+: https://scenicplus.readthedocs.io/en/latest/.

SCENIC+ is a three-step workflow consisting of: Identifying candidate enhancers; Identifying enriched TF binding motifs on candidate enhancers; Linking TFs to candidate enhancers and target genes, as shown in Figure 4A.

#### 1. Identifying candidate enhancers

SCENIC+ uses both differentially accessible regions (DARs) and topics (sets of co-accessible regions) across cell types or states as enhancer candidates. In this work, for each of 95 cell states, we identified DARs using ArchR (FDR<0.01, Log2FC>0.5), which resulted in a total of 122,554 DARs. In addition, the topic modeling was performed with pycisTopic using Latent Dirichlet Allocation (LDA) with the collapsed Gibbs sampler to iteratively optimize the probability of a region belonging to a topic. A model of 200 topics was selected based on the stabilization of metrics described in the references and log-likelihood. Region–topic probabilities were binarized using either the Otsu method or by selecting the top-3,000 regions per topic. These DARs and region–topic associations served as the starting point for further analyses to identify enhancer–gene links and eGRNs.

#### 2. identifying enriched TF-binding motifs on candidate enhancers (cistrome)

To discover potential transcription factor binding sites (TFBSs) in candidate enhancers, we conducted motif enrichment analysis. We started by collecting 590 motifs from DANIO-CODE(Baranasic et al., 2022), corresponding to 912 transcription factors (TFs), since some TFs share the same motif. We used motifMatch(Schep et al., 2017) to create a motif match score for each motif in each candidate enhancer (region). We then performed motif enrichment analysis using pycisTarget. Motif enrichment was conducted with both the cisTarget and differential enrichment of motifs (DEM) algorithms on cell-type-based DARs, the top 3,000 regions per topic, and topics binarized using the Otsu method. cisTarget is a ranking-and-recovery-based algorithm, where enrichment is calculated as a normalized area under the curve (AUC) at the top 0.5% ranking, resulting in a normalized enrichment score (NES). Motifs with an NES greater than 3.0 were retained. To identify the target regions for each motif (motif-based cistrome), regions at the top of the ranking (leading edge) were retained, defined by an automated thresholding method that keeps regions below the rank at maximum enrichment. For the DEM algorithm, a Wilcoxon rank-sum test was performed between a foreground and a background set of regions using score distributions for each motif or cluster of motifs. Motifs with an adjusted p-value less than 0.05 (Bonferroni) and log fold change greater than 0.5 were kept. Regions containing the motif (motif-based cistrome) were obtained by selecting regions with a cis-regulatory module score greater than 3 for each enriched motif.

#### 3. linking TFs to candidate enhancers and target genes (eRegulon)

##### 3.1 Calculating region-to-gene and TF-to-gene importance scores

We used the default parameters from SCENIC+ to quantify region-to-gene importance scores. In SCENIC+, the importance score for each region was calculated using gradient-boosting machine regression to predict target gene expression based on region accessibility. All regions within a gene’s search space, defined as a minimum of 1 kb and a maximum of 150 kb upstream or downstream of the gene’s start or end, or the promoter of the nearest upstream or downstream gene, were considered.

The promoter of a gene was defined as the transcription start site ±10 bp. Since the importance score does not indicate directionality, we calculated the Spearman rank correlation between accessibility and expression, using the correlation coefficient to distinguish positive interactions (>0.03) from negative interactions (<−0.03). Similarly, for each TF and gene pair, we calculated the importance score of TF expression to predict target gene expression using gradient-boosting machine regression, and used Pearson correlation to separate positive (>0.03) from negative (<−0.03) interactions.

##### 3.2 Binarizing region-to-gene importance scores

We used the default parameters from SCENIC+ to binarize region-to-gene importance scores. In SCENIC+, region-to-gene importance scores were binarized using multiple methods: taking the 85th, 90th, and 95th quantiles of the importance scores, selecting the top 5, 10, and 15 regions per gene based on these scores, and applying a custom implementation of the BASC88 method on the region-to-gene importance scores.

##### 3.3 eRegulon creation

We used the default parameters from SCENIC+ to create eRegulons based on the cistrome, the region-to-gene, and the TF-to-gene relationships. For each TF, TF–region–gene triplets were identified by first selecting all regions enriched for motifs associated with the TF, based on cistrome data. Next, we used binarized region-to-gene links to assign genes to these regions. To eliminate false positives, we conducted gene set enrichment analysis (GSEA), ranking genes by their TF-to-gene importance score and calculating the enrichment of the gene set within the TF–region–gene triplet using the gsea_compute function from GSEApy. Genes at the top of the ranking (the leading edge) were retained as target genes of the eRegulon. eRegulons with fewer than ten predicted target genes were discarded.

#### 4. eRegulon activity

All consensus peaks and genes were ranked based on their chromatin accessibility and raw gene expression counts per cell, respectively. Enrichment for eRegulon target regions and target genes was defined as the Area Under the Curve (AUC) at the top 5% of the ranking. This enrichment score is defined as eRegulon activity, as shown in Figure 4C, D and Table S5. There are two types of eRegulon activity: one based on target regions and the other based on target genes.

#### 5. eRegulon filter

eRegulons were filtered based on the correlation coefficient between the AUC scores (eRegulon activity) of the target regions and the target genes. eRegulons with a correlation coefficient greater than 0.4 were considered high quality. For downstream analysis, we further refined the selection to include only activator eRegulons, defined as those with both region-to-gene and TF-to-gene correlations greater than 0. This filtering process resulted in 100 eRegulons (as shown in Figure 4B), with a median of 70 genes and 93 regions per eRegulon.

#### 6. eRegulon specificity score

For each eRegulon, we calculated an eRegulon specificity score (RSS, as shown in Figure 4C and Table S6) in each of the 95 cell states using Jensen-Shannon divergence, which measures the similarity between two probability distributions. This calculation utilized the target genes based on eRegulon activity as input. We determined the Jensen-Shannon divergence by comparing each vector of binary eRegulon activity overlaps with the assignment of cells to specific cell types.

### Mis-expression

#### 1. Cloning and vector construction

For EVL TFs, open reading frames (ORFs) were PCR-amplified with Platinum SuperFi II DNA Polymerase (Invitrogen) using primers with overhanging restriction sites (BstI for forward primers, SpeI or AgeI for reverse primers). PCR products were purified using MinElute PCR Purification Kit (QIAGEN) and digested using BstI and SpeI or BstI and AgeI (New England Biolabs) at 37°C for 1 hour. The digested amplicons were then purified using MinElute PCR Purification Kit (QIAGEN). An in vitro transcription vector containing a beta-globin 5′ UTR sequence with an optimized translation initiation site was used for optimal mRNA stability (https://doi.org/10.1101/2023.11.23.568470). This vector was digested using BstI and AgeI or BstI and SpeI (New England Biolabs, NEB) at 37°C for 1 hour, dephosphorylated using Antarctic Phosphatase (New England Biolabs #M0289) at 37°C for 30 minutes, followed by purification using MinElute PCR Purification Kit (QIAGEN). The ORF amplicon was directionally ligated to the vector using T4 DNA ligase (New England Biolabs #M0202S) and transformed into One Shot™ TOP10 Chemically Competent E. coli cells (Invitrogen) using the heat shock method. Plasmid DNA was isolated from these cultures using QIAprep Spin Miniprep Kit (QIAGEN). The resulting constructs were validated by Sanger sequencing. For YSL TFs, synthetic gene fragments with flanking BstI and SpeI/AgeI restriction sites were ordered from Twist Bioscience for cloning.

#### 2. mRNA synthesis and purification

Validated constructs were used as templates for PCR amplification with Platinum SuperFi II DNA Polymerase (Invitrogen) using an SP6 forward primer (5′ CACGCATCTGGAATAAGGAAGTGC 3′) and a 3′ UTR-specific reverse primer with a 36 nt-long poly(T) overhang (5′ TTTTTTTTTTTTTTTTTTTTTTTTTTTTTTTTTTTTTCCTGTGAGTCCCATGGGTTTAAG 3′) as described in (https://doi.org/10.1101/2023.11.23.568470). PCR products were purified and transcribed using mMESSAGE mMACHINE™ SP6 Transcription Kit (Invitrogen). The resulting mRNA was purified using RNA Clean & Concentrator™-5 (Zymo Research).

#### 3. mRNA misexpression and embryo dissociation

Embryos were obtained from wild-type (Tupfel longfin/AB) and microinjected with 15 pg of each TF mRNA (total of ∼100 pg) at the one-cell stage. This was determined to be the optimal dose with minimal embryo lethality at 50% epiboly. Injected embryos were incubated at 28.5°C in embryo medium. At approximately 30% epiboly, ∼50 embryos per experimental condition were dechorionated using 1 mg/ml Pronase (Protease type XIV from Streptomyces griseus, Millipore Sigma). At 50% epiboly, dechorionated embryos were added to a protease solution (10 mg/ml BI protease [Sigma, P5380], 125 U/ml DNaseI, 2.5 mM EDTA in DPBS) in 2 ml LoBind Eppendorf tube (EP-022431048) and incubated on ice for 4 minutes followed by the addition of a stop solution stop solution (30% FBS, 0.875 mM CaCl2 in DPBS). The supernatant was removed and replaced with chilled Ringer’s solution (140 mM NaCl, 2 mM KCl, 1.5 mM K2HPO4, 1 mM MgSO4, 2 mM MgCl2, 10 mM HEPES, 10 mM D+ glucose) followed by gentle pipetting (5-10 times) to dissociate the embryos. The cells were pelleted at 4°C with 250g for 4 minutes to discard the supernatant. This dissociation process was repeated two more times with chilled Ringer’s solution followed by two more times with chilled DMEM (Gibco, 11594426) with 0.1% BSA followed by resuspension in a final volume of 500 μl. The cells were filtered through a 70 μm cell strainer (Flowmi Cell Strainer, BAH136800070) into 15 mL polypropylene Falcon tubes (Corning 352196) pre-coated with 1% BSA. Cell viability and concentration were evaluated using AO-PI viability dye (0.002% acridine orange and 0.02% propidium iodide) on a hemocytometer.

#### 4. Single-cell RNA sequencing using Parse Bioscience

Single-cell suspensions were immediately fixed using Evercode™ WT v2 (Parse Biosciences) according to the manufacturer’s protocol. Approximately 4000-5000 cells per condition were used for library construction using either Evercode™ WT Mini v2 or Evercode™ WT v2 (Parse Biosciences). Two sub-libraries were generated using Evercode™ WT Mini v2 and eight sub-libraries using Evercode™ WT v2.

All sub-libraries were sequenced on a NovaSeq platform (Illumina).

#### 5. Data analysis

The resulting FASTQ files were demultiplexed using the *split-pipe* pipeline (v1.2.1) from Parse Biosciences and aligned to Ensembl GRCz11 reference genome (Ensembl Release 100). The data was processed and analyzed using Seurat version 4.1.0(Hao et al., 2021). Cells with fewer than 1,000 expressed genes or more than 6,000 expressed genes were filtered out. On average, we obtained 3,466 cells per sample, with each cell expressing an average of 3,070 genes (Table S8). Each sample was processed separately, with cell types clustered and annotated accordingly.

### Data access

The processed data are saved in: https://drive.google.com/drive/folders/1NTRCoIOviDVmsVhDiaPXpbdDqs40D7rz?usp=drive_link The folder includes the following files:

- Seurat object of the nine-stage integrated single-cell multi-omic datasets
- Peak file
- Single-cell metadata file
- eRegulon file
- eRegulon specificity file
- DeepDanio models and example code for predicting chromatin accessibility based on DNA sequences
- DeepDanio identified motifs and TFBSs for each of the 95 cell states

## Acknowledgements

We thank the Schier lab for input and discussions. We thank Alba Aparicio Fernandez, Rita Gonzalez Dominguez and Diana Medeiros Gomes for support with fish husbandry. We thank Fabien Cubizolles for support with experimental work. High-throughput sequencing was performed at the Genomics Facility Basel. Part of the computations were performed at the sciCORE Center for scientific computing at the University of Basel. This work was funded by ERC Advanced grant 834788 and the Allen Discovery Center for Cell Lineage Tracing to A.F.S., NIH Award R33CA255893 and NSF Award 2021552 to G. S.

## Contributions

J.L., and A.F.S. conceived and designed the study. J.L., Y.W., and A.N.C. collected single-cell multi-omics data. J.L. performed single-cell multi-omics related analyses. S.M.C-H. designed and trained deep learning model. S.M.C-H., J.L., and C.Y. performed deep learning related analyses. J.L. reconstructed GRN and performed related analyses. L.Y.D. led the mis-expression experiments with help from J.L., and M.C-R. J.L. analyzed the mis-expression data. J.L., A.F.S., L.Y.D., S.M.C-H., M.C-R. C.Y. and G.S. interpreted the results. J.L. wrote the original draft, J.L., A.F.S., G.S. L.Y.D., and S.M.C-H. finalized the paper with input from all the other authors. All authors read and approved the final manuscript.

---

**Figure S1:**
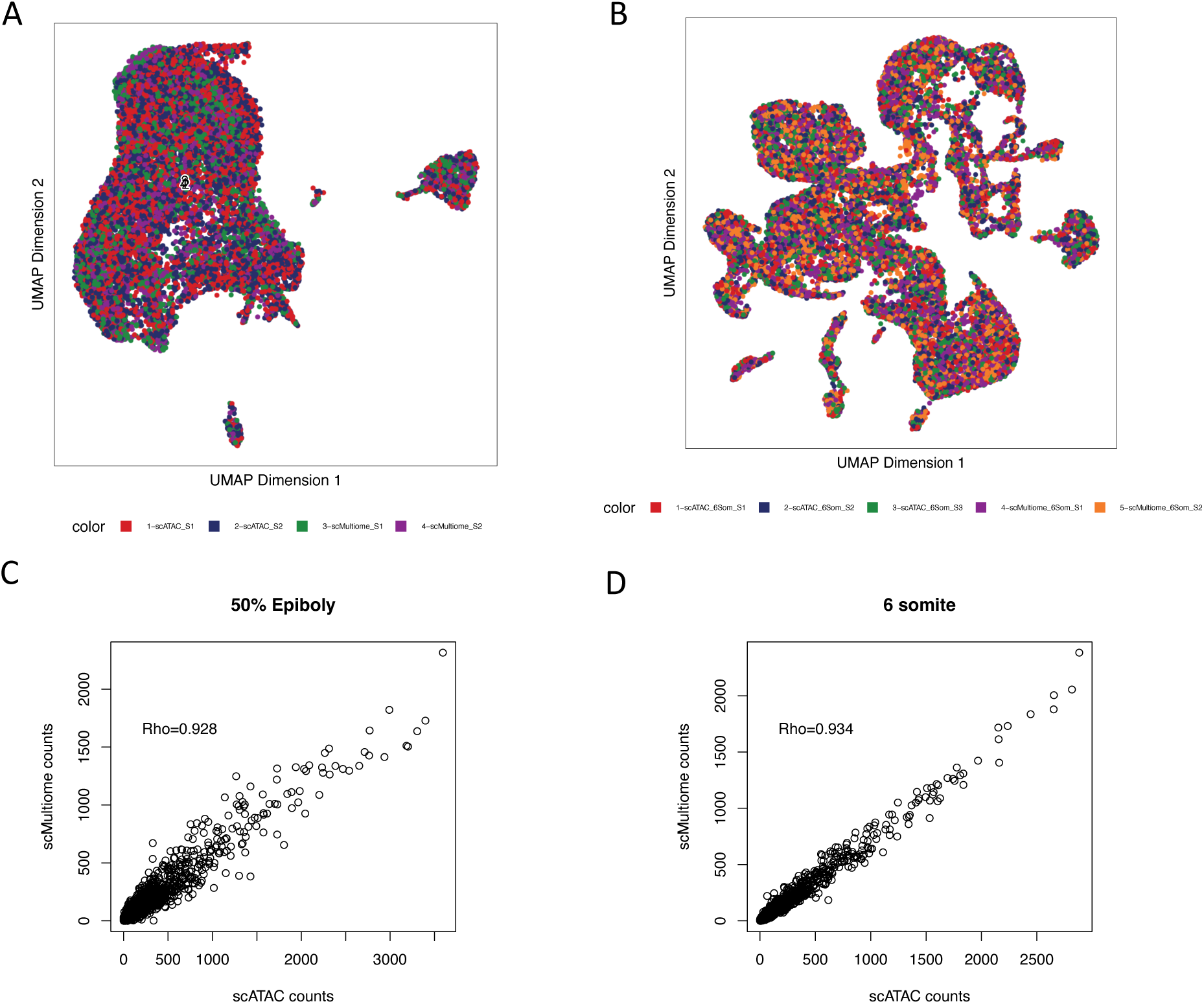
Comparison of scMultiome ATAC and standalone scATAC. A and B. UMAP visualization of single cells, with each cell colored according to its respective condition. S1, S2, and S3 represent different technical replicates. C and D. Count correlation (spearman’s rank correlation) between scMultiome ATAC and standalone scATAC. The genome was divided into 500 bp bins, and the fragment counts were calculated for each bin in each cell. These counts were then aggregated across single cells to create a pseudobulk-count. The correlation was performed on these pseudobulk-counts.

**Figure S2:**
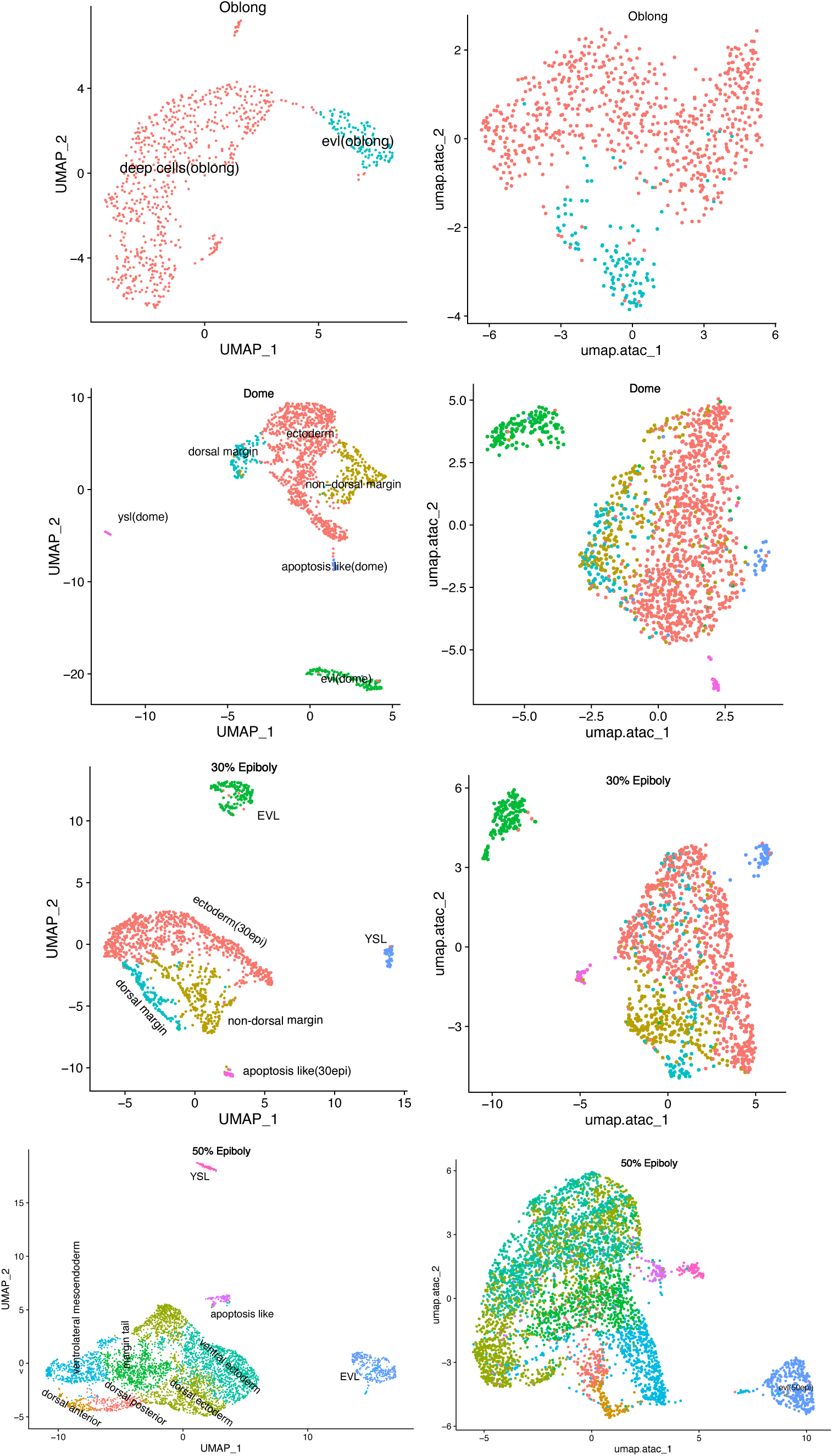
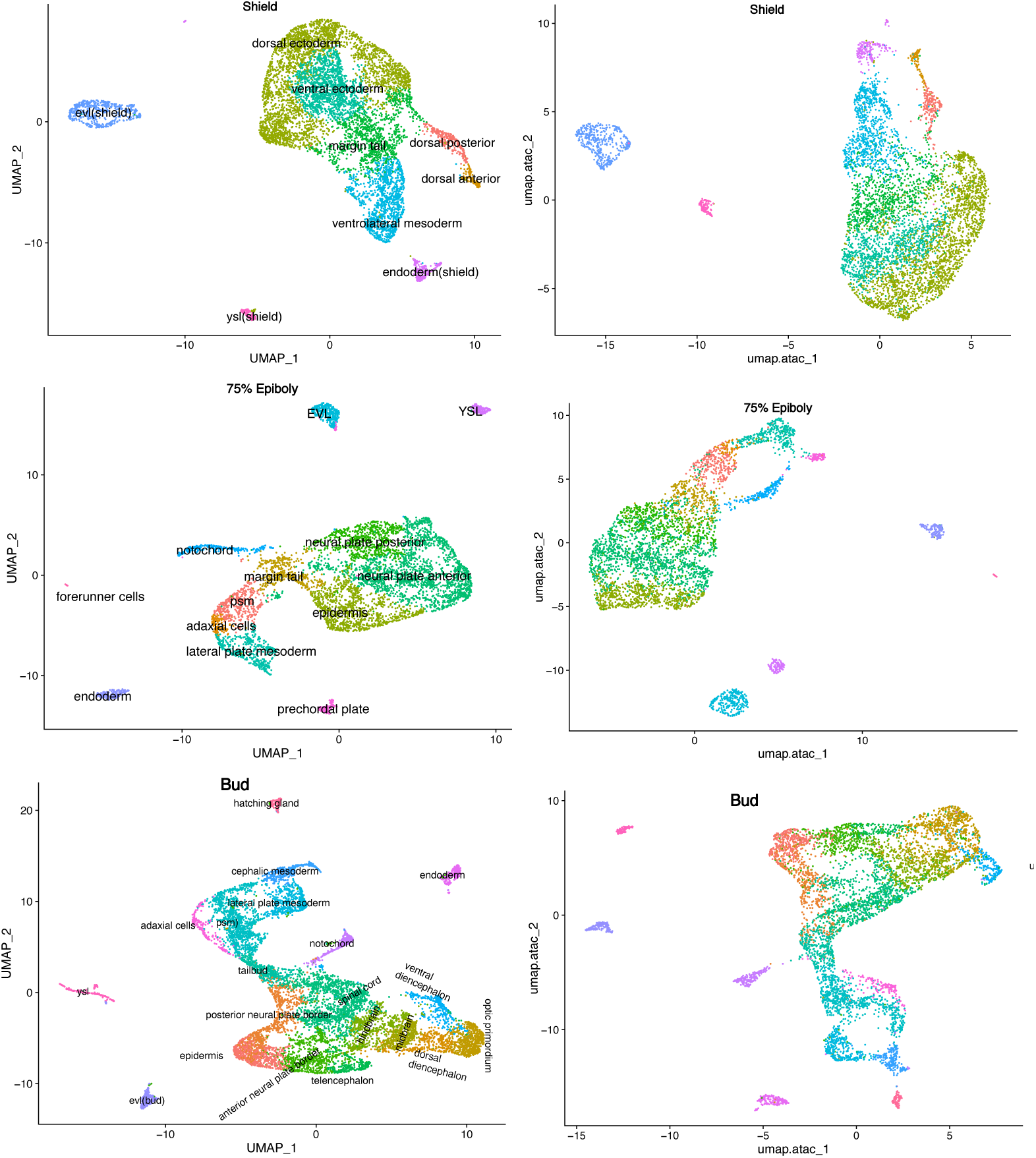
UMAP visualization of cell state diversification from the oblong to bud stages. The clusters/cell states are defined by gene expression, and the UMAP coordinates derived from either RNA-seq data (left) or ATAC-seq data (right).

**Figure S3:**
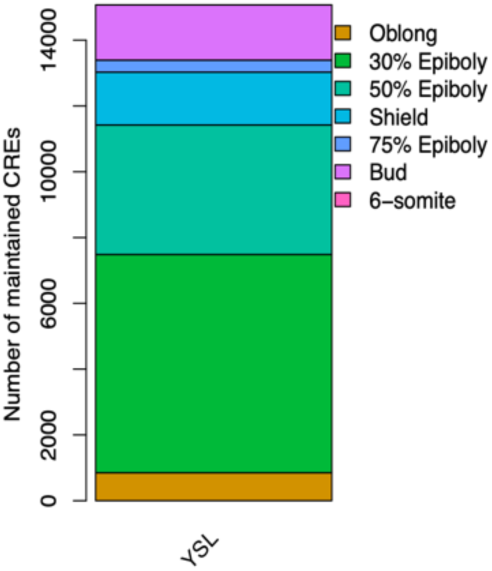
The number of acquired dCREs at each stage that are maintained by the 6-somite stage for the YSL trajectory. Due to the low number of YSL cells at the dome stage (only 22 cells), this stage was excluded from the analysis.

**Figure S4:**
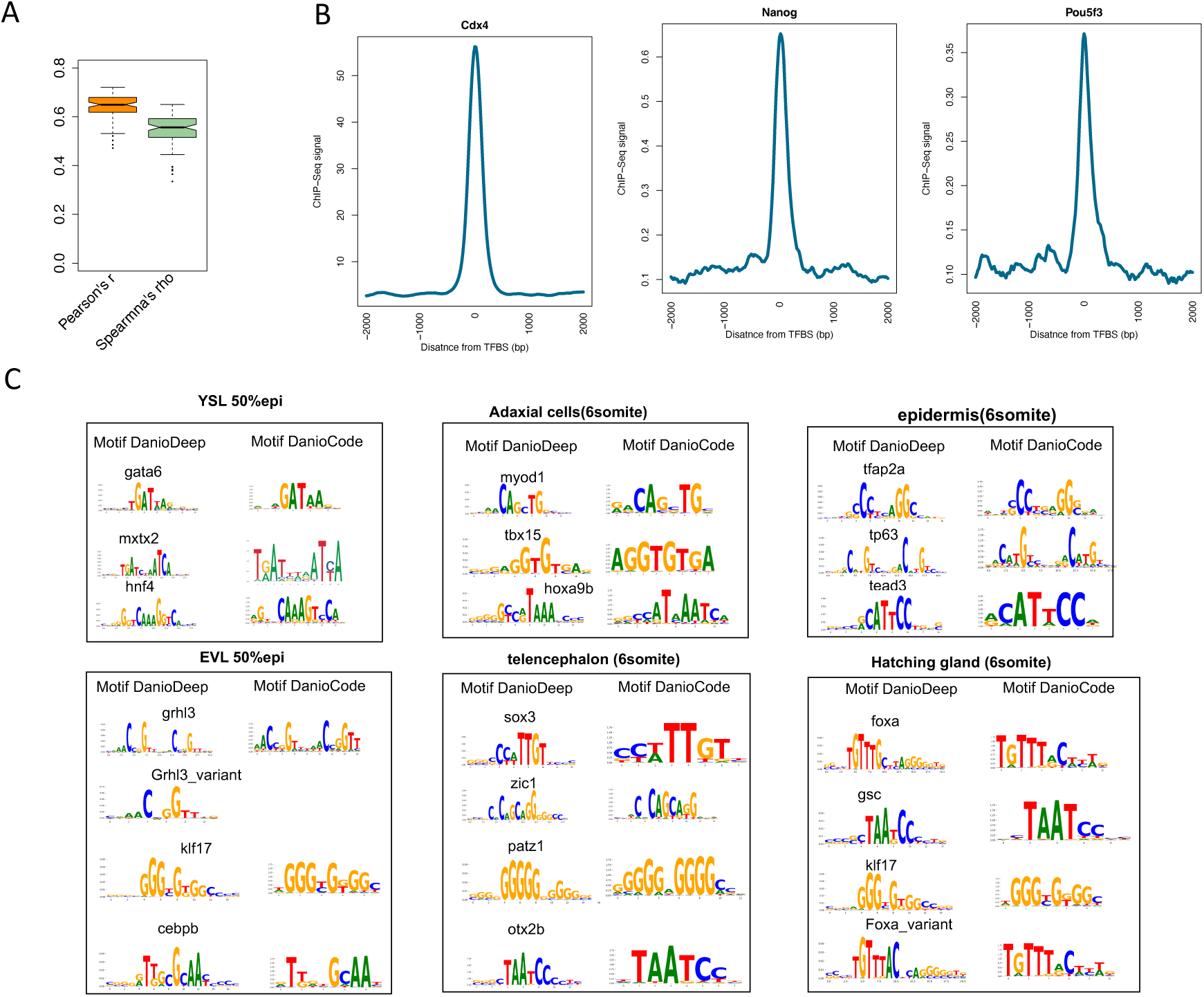
Deep learning quality control. A. For each cell type, we calculated the Pearson or Spearman correlation coefficient between the predicted and actual chromatin accessibility. The boxplot shows these correlations, with the lower and upper whiskers representing 1.5 times the interquartile range (IQR). The box itself displays the IQR, with the median indicated. B. ChIP-seq signal in either 2kb upstream or downstram of TFBS. C. De novo motifs identified by DeepDanio were annotated using motifs from the DANIO-CODE database.

**Figure S5:**
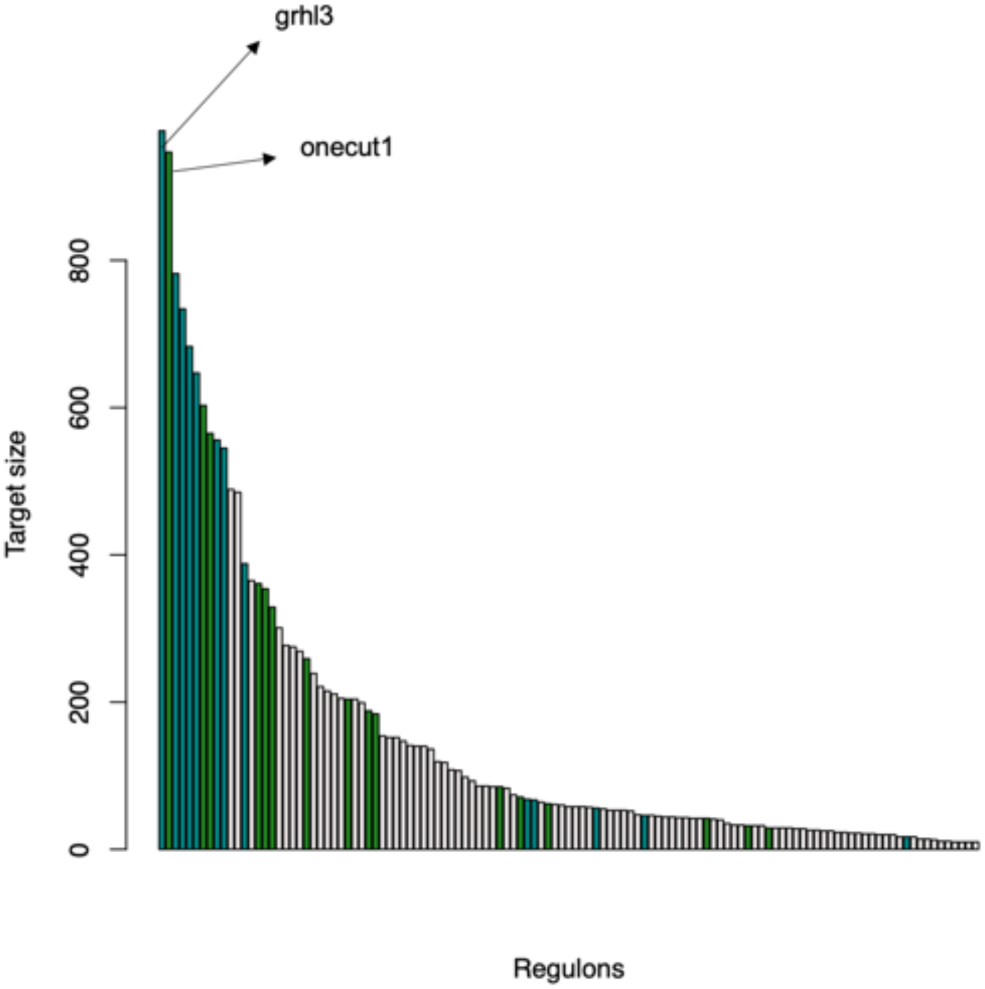
Histogram showing the number of genes regulated by each TF. Blue represents EVL eRegulons, while green represents YSL eRegulons. The top 10 specific eRegulons in each cell state of the EVL (from oblong to 6-somite) and YSL (from dome to 6-somite) were selected, combined, and filtered to create unique sets of EVL and YSL eRegulons.

**Figure S6:**
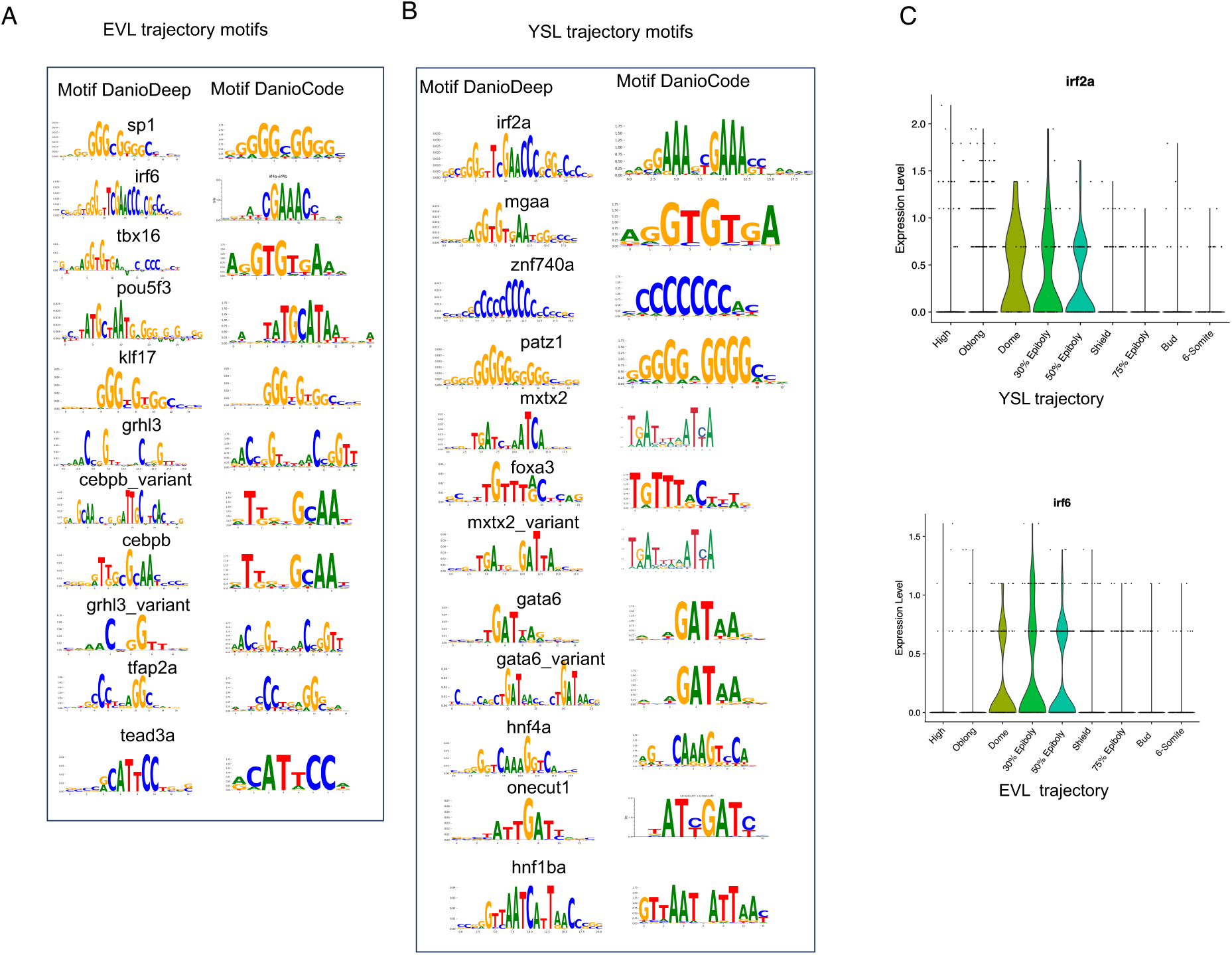
De novo motifs identified by DeepDanio were annotated using motifs from the DANIO-CODE database. A. De novo motifs identified by DeepDanio for the EVL trajectory. For each cell state along the EVL trajectory, the top 5 motifs were selected. Since some motifs were shared between stages, all top 5 motifs from each cell state were combined and filtered to obtain a unique set of motifs. This figure displays the resulting unique set of motifs for the EVL trajectory. B. De novo motifs identified by DeepDanio for the YSL trajectory. For each cell state along the YSL trajectory, the top 5 motifs were selected. Since some motifs were shared between stages, all top 5 motifs from each cell state were combined and filtered to obtain a unique set of motifs. This figure displays the resulting unique set of motifs for the YSL trajectory. C. Expression of irf2a and irf6. The same motif was assigned to different transcription factors based on expression levels. The second motif in EVL was annotated as irf6 due to irf6 being expressed in EVL, while the first motif in YSL was annotated as irf2a because irf2a was expressed in YSL.

**Figure S7:**
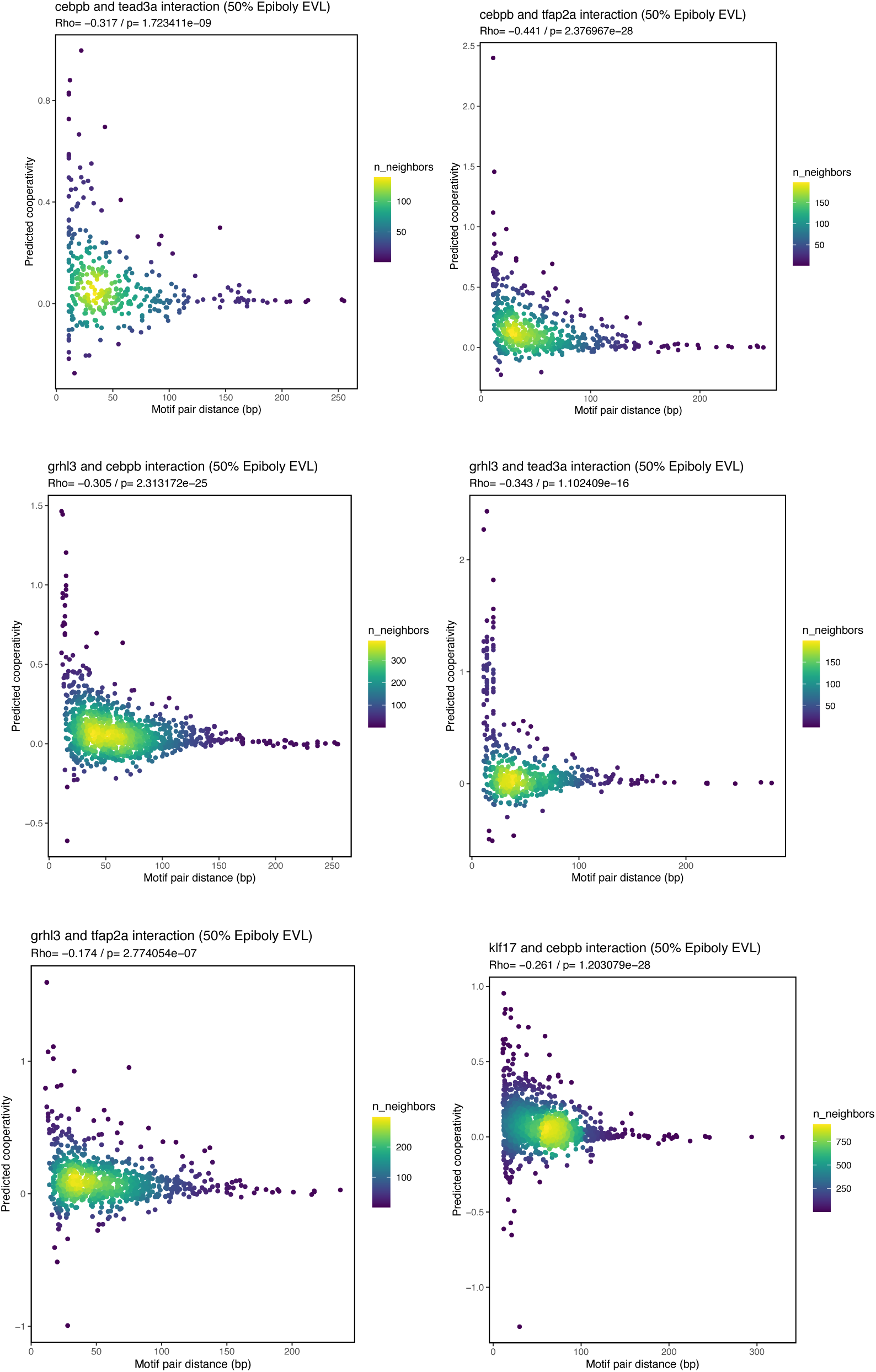
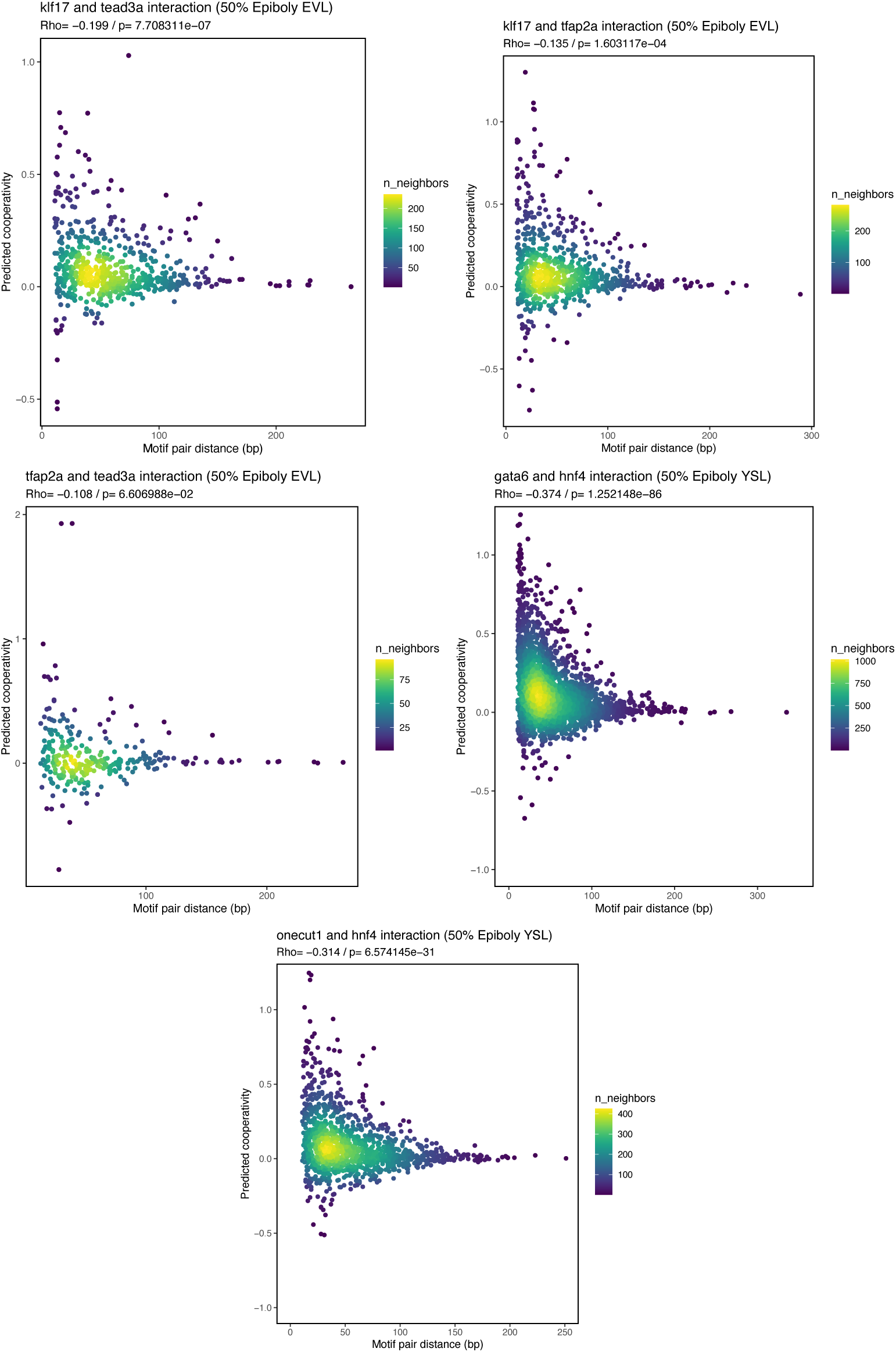
Correlation between distances of two TFBSs and interaction scores. The Spearman’s correlation coefficient (Rho) and the P-value are displayed in the top-left corner of the plot.

**Figure S8:**
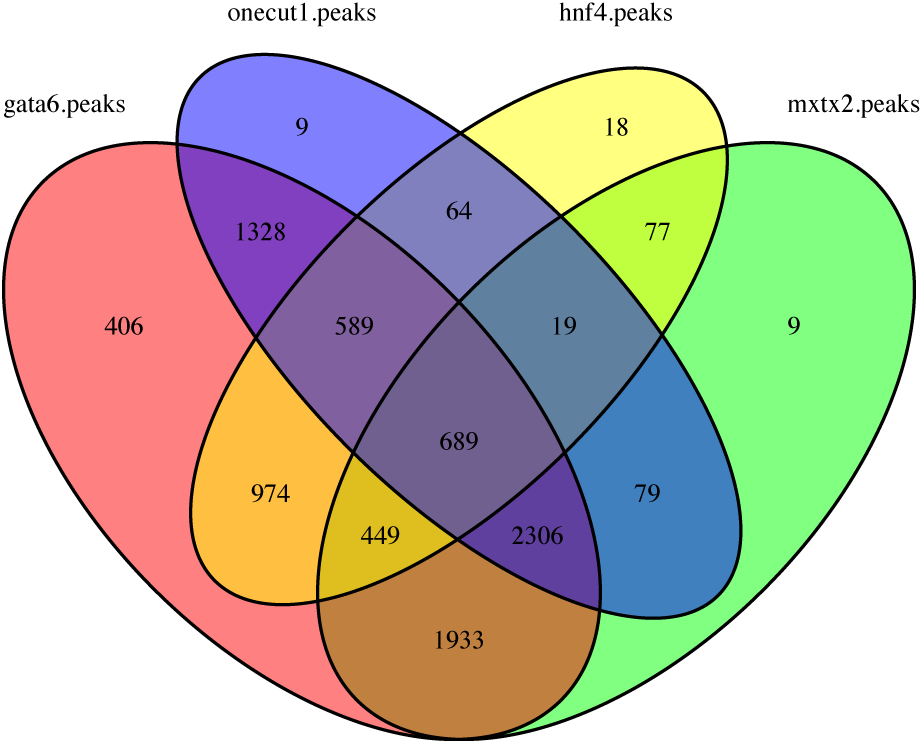
Venn Diagrams of CREs containing gata6, hnf4, onecut1, and mxtx2 Motifs.

**Figure S9:**
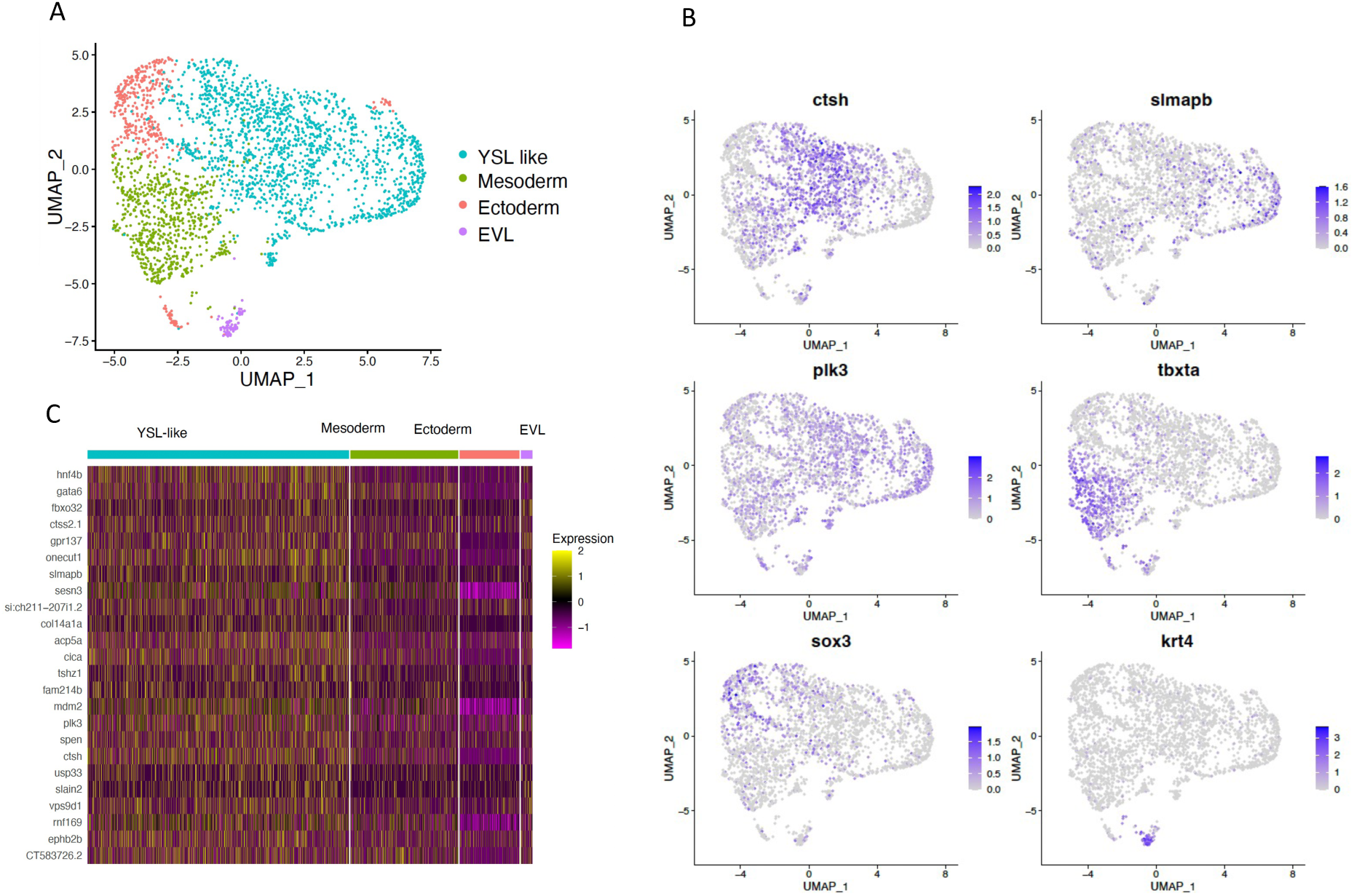
YSL misexpression without mxtx2. A. UMAP projection of single-cell transcriptomes, with cells colored by cell type. B. UMAP projection of single-cell transcriptomes, with cells colored by marker gene expression. *ctsh*, *slmapb*, and *plk3* are YSL marker genes, *tbxta* is a marker for mesoderm, *sox3* is a marker for ectoderm, and *krt4* is a marker for EVL. C. Heatmap showing the expression of YSL-specific genes that are activated by the misexpression of all differentiation TFs, excluding Mxtx2.

**Figure S10:**
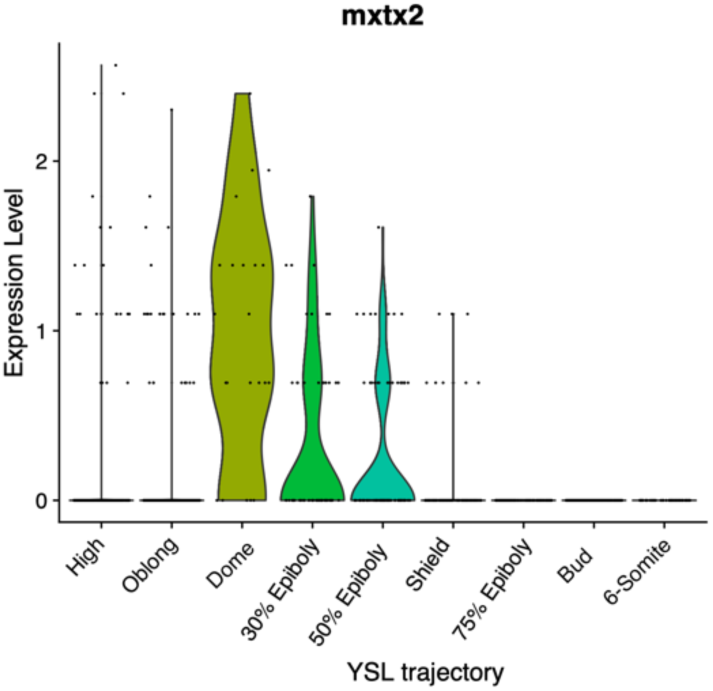
*mxtx2* expression across YSL trajectory.

**Figure S11:**
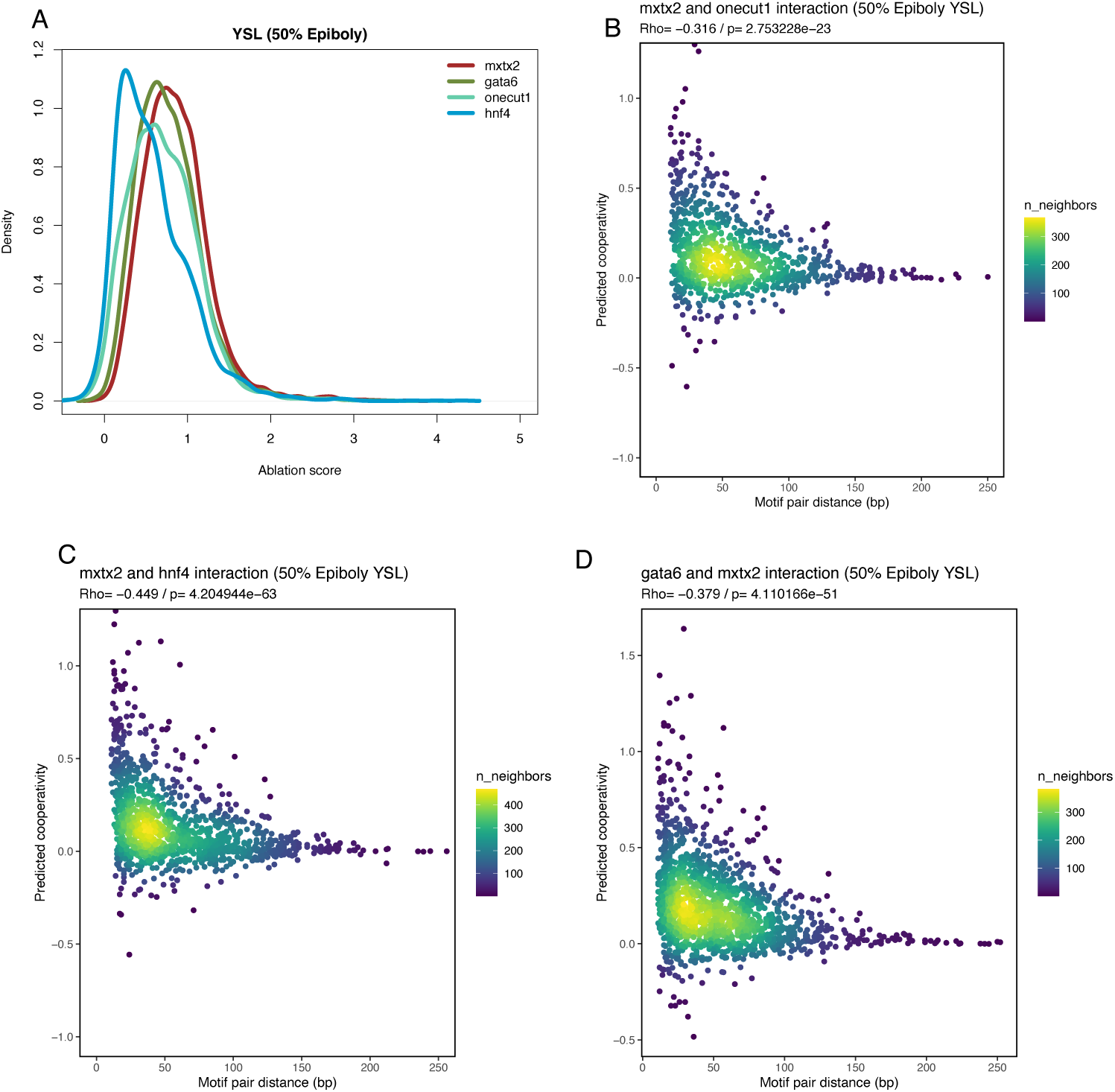
ablation analysis including mxtx2 motif. A: Distribution of ablation scores for each TF B, C, D: Correlation between distances of two TFBSs and interaction scores. The Spearman’s correlation coefficient (Rho) and the P-value are displayed in the top-left corner of the plot.

